# Environmental Conflict Modulates Pavlovian Bias

**DOI:** 10.64898/2025.12.22.696027

**Authors:** Priyanshu Priyanshu, Arjun Ramakrishnan

## Abstract

Pavlovian avoidance enables rapid defensive responding but can undermine goal-directed behaviour when it overrides instrumental control, a tendency amplified in anxiety. Whether such biases can be flexibly and rapidly modulated by environmental structure remains unknown. Here we show that global conflict in the environment can suppress Pavlovian avoidance and enhance instrumental choice, particularly in individuals high in trait anxiety. One hundred and sixteen participants completed a sequential approach–avoidance task in which three options differed in reward–punishment conflict, and blocks varied the frequency of high-conflict encounters. In low-conflict environments, participants exhibited pronounced and misplaced avoidance of objectively safe, no-conflict options indicating that choices are not determined by local contingencies alone but are tuned to broader environmental context. Interestingly, increasing environmental conflict reduced this maladaptive avoidance: approach to safe options increased, and reinforcement-learning and drift–diffusion modelling revealed higher reward sensitivity and drift rates alongside reduced lapse and Pavlovian bias parameters. These effects were strongest in high-anxious individuals, whose behaviour shifted markedly toward that of low-anxious peers. Crucially, benefits depended on the frequency of conflict encounters and were not elicited by cue–outcome reversals or increased punishment probability alone. Furthermore, a modified Go/NoGo task replicated global conflict-dependent improvements on Pavlovian-incongruent trials. Together, these results demonstrate that global environmental conflict can rapidly recalibrate the balance between Pavlovian and instrumental systems, attenuating maladaptive avoidance and disproportionately aiding anxious individuals. We propose that conflict-driven arousal dynamics, plausibly mediated by LC–NE mechanisms, promote this adaptive reconfiguration of defensive and goal-directed control.

**Significance statement:** Avoiding danger is essential for survival, but sometimes our automatic fear responses make us avoid things that are actually safe and potentially rewarding. This problem is especially common in people with anxiety, where avoidance can become rigid and limit everyday functioning. In this study, we discovered that the broader structure of the environment, not just the immediate situation, can strongly influence these automatic reactions. When the environment contained more frequent conflicts or difficult choices, people actually became less avoidant, even toward options that were completely safe. Their decisions became more goal-directed, and this shift was especially strong in individuals with high anxiety, whose behaviour began to resemble that of low-anxiety individuals. These results show that anxious individuals can be more flexible and their behaviour can be rapidly improved by changing the surrounding context. This challenges assumptions about how rigid avoidance tendencies are and suggests ways to reduce maladaptive avoidance without relying solely on long or intensive training. By revealing how environmental structure can rebalance automatic fear responses and deliberate decision-making, this work opens avenues for designing interventions and potentially therapies that harness contextual factors to support more adaptive behaviour in anxiety.

## Introduction

Learning through association enables humans to rapidly extract predictive structure from their environment and deploy adaptive responses with minimal deliberation (Bromiley, 1949; Gámez et al., 2013; Pavlov, 2010; Pontes et al., 2020; Shanks, 2010). Pavlovian learning promotes reflexive approach toward rewards and avoidance of punishments, whereas instrumental learning supports slower but flexible, goal-directed action based on action–outcome contingencies (Dayan & Balleine, 2002; Gershman, 2015; Raab & Hartley, 2020). Although these systems are computationally distinct, human behaviour reflects their continuous interaction. When aligned, they support efficient learning; when misaligned, automatic Pavlovian tendencies can override instrumental goals, producing impulsive or avoidant responses that undermine adaptive choice (Algermissen & Den Ouden, 2023; Gershman et al., 2021; Pool et al., 2019; Saeedpour et al., 2023; Weber et al., 2022).

Pavlovian biases are particularly consequential in situations involving mixed incentives, where approach–avoidance conflicts arise (Bublatzky et al., 2017; Guitart-Masip et al., 2012; Hegefeld & Davidow, 2025; Huys et al., 2011). Excessive avoidance in such contexts incurs significant opportunity costs (Wright et al., 2013; Pittig & Scherbaum, 2020; Boschet et al., 2022; Sayalı et al., 2025) and is a hallmark of anxiety and related psychopathologies (Huys et al., 2016; Nord et al., 2018; Eisinger et al., 2020; Peterburs et al., 2022; Fleming et al., 2023; Goldman et al., 2024; Hulsman et al., 2024; Burghoorn et al., 2025; Embrey et al., 2025). While recent work shows that Pavlovian biases can be attenuated through semantic framing or extended training (Ereira et al., 2021; Fleming et al., 2023; Moutoussis et al., 2018), it remains unknown whether these biases can be modulated *rapidly* and *adaptively* in response to contextual demands, and how such modulation differs as a function of anxiety.

Overriding prepotent avoidance to pursue long-term goals requires flexible engagement of instrumental control (Dorfman & Gershman, 2019; Mahajan et al., 2025; Millner et al., 2018). A key candidate mechanism for enabling such flexibility is the locus coeruleus–norepinephrine (LC-NE) system, which regulates arousal and attentional mode through tonic and phasic dynamics (Aston-Jones & Cohen, 2005; Howells et al., 2012; De Gee et al., 2017; Breton-Provencher et al., 2021; Grueschow et al., 2021). These modes govern shifts between focused exploitation and broader environmental monitoring and play a critical role in flexible decision-making across reinforcement-learning contexts (Aston-Jones & Cohen, 2005; Jepma, 2010; Vazey et al., 2018; Van Der Linden et al., 2021; Grimm et al., 2024). Dysregulation of LC-NE activity is implicated in the heightened rigidity and threat sensitivity characteristic of anxiety (McCall et al., 2017; Morris et al., 2020; Jain et al., 2024; Boukezzi et al., 2025).

Building on this framework, we propose that LC-NE dynamics regulate transitions between Pavlovian and instrumental systems. Specifically, we hypothesize that tonic NE tracks global environmental conflict or volatility, whereas phasic NE responds to trial-level conflict. Drawing on the Yerkes–Dodson principle, we predict that increasing environmental conflict elevates tonic NE toward an optimal arousal zone that facilitates instrumental control and suppresses prepotent Pavlovian responding. Because high-trait-anxious individuals operate at elevated baseline arousal, even modest increases in conflict may shift them more efficiently into this optimal state, predicting greater conflict-induced reductions in Pavlovian bias and a crossover pattern of behavioural flexibility.

These considerations motivate our central question: Does environmental conflict, via LC-NE–mediated arousal shifts, enable rapid suppression of Pavlovian bias, and does trait anxiety modulate this conflict-dependent rebalancing between Pavlovian and instrumental control?

Addressing this question requires moving beyond classic paradigms such as Go/No-Go, probabilistic reversal learning, and Pavlovian-instrumental transfer tasks, which have been significant in revealing Pavlovian constraints on instrumental control under punishment or threat. However, these tasks often rely on simplified and weakly ecological action–outcome mappings such as arbitrary cues requiring action suppression for reward or action execution to avoid punishment that depart from the natural statistics of real-world approach–avoidance decisions. Moreover, they typically probe control failures at the level of individual trials or static contingencies, offering limited leverage on how broader environmental structure such as the global frequency of conflict shapes the balance between Pavlovian and instrumental systems over time.

To address this gap, we developed a novel approach–avoidance paradigm that parametrically manipulated environmental conflict, contingency reversals, and aversive contexts while engaging the LC-NE system. Participants displayed generalized, misplaced avoidance even under no-conflict conditions, but increasing conflict robustly suppressed Pavlovian bias. Moreover, individuals high in trait anxiety exhibited both heightened baseline bias and the strongest conflict-induced attenuation. Importantly, we observed a convergent pattern in a modified Go/No-Go task in which global conflict was similarly manipulated: increasing environmental conflict reduced Pavlovian bias in this classic paradigm as well, indicating that conflict-dependent recalibration generalizes beyond task-specific structure. Together, these findings outline a neurocomputational account of how anxiety and environmental structure jointly shape human decision-making by reconfiguring the balance between reactive and goal-directed systems.

## Results

One hundred and sixteen participants completed a sequential approach–avoidance task in which, on every trial, they encountered a tree they could either approach or avoid. Approaching offered the possibility of reward but also carried the risk of receiving an electric shock paired with monetary loss, thereby inducing an approach–avoidance conflict (Fig. 1a,b). There were three types of trees that varied in levels of conflict they induced: a no-conflict tree (NCT) which provided 1 unit of Reward or No Punishment, a low-conflict tree (LCT) that provided 2 units of reward or one unit of punishment, and a high-conflict tree (HCT) that provided 3 units of reward or 2 units of punishment upon approaching (Fig. 1b). Furthermore, by altering the relative frequency with which these trees appeared across blocks, environmental/ global conflict was manipulated. In the low-conflict environment (LCE), participants predominantly encountered NCTs (45%) and rarely HCTs (20%). In the subsequent high-conflict environment (HCE), this distribution was reversed, exposing participants more frequently to high-conflict choice (Fig 1c).

**Figure 1:**
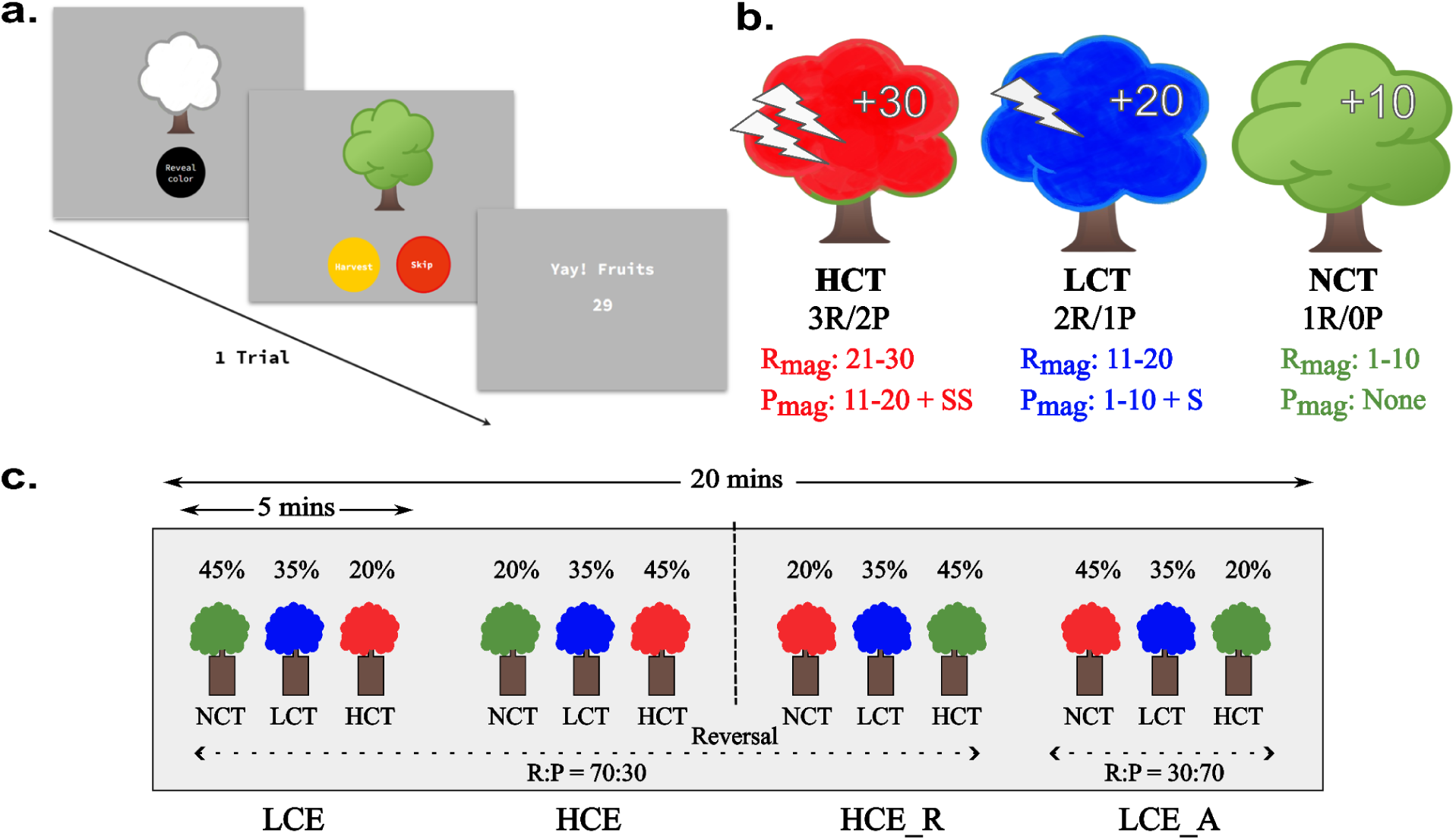
Approach avoidance conflict paradigm. a. Trial structure of the task, white tree with reveal button followed by a colored tree (Red/ Blue/Green) with options to approach or avoid, finally the feedback (Reward or Punishment). b. Red, Blue and Green colored trees representing High (HCT), Low (LCT) and No (NCT) conflict stimulus with their respective reward-punishment offers,a high-conflict tree (HCT) that provided 3 units of reward (R) or 2 units of punishment (P) and shock, a low-conflict tree (LCT) that provided 2 units of reward and one unit of punishment and shock, and a no-conflict tree (NCT) which provided 1 unit of Reward and No Punishment upon approaching with actual ranges of Reward (R_mag_) and Punishment (P_mag_) magnitudes given below. c. Task orientation with a single stretch of continuous design lasting 20 minutes, however, consists of 4 blocks of 5 minutes each, first, low conflict environment(LCE) with 45% NC, 35% LC and 20% HC trees. Second, a high conflict environment (HCE) with 20% NC, 35% LC and 45% HC trees. Third, HCE with reversal (HCE_R), contingencies of HC and NC were reversed i.e. color between HCT and NCT were reversed breaking cue-associations. For the first three blocks the reward: punishment ratio was maintained at 70:30 respectively but it was reversed for fourth block i.e. LCE with aversiveness (LCE_A) flipped reward-punishment to 30:70.

### Approach–avoidance decisions reflect reward–punishment conflict and Pavlovian influence

We first examined the Low Conflict Environment (LCE) and asked whether choice behaviour and reaction times (RTs) were sensitive to conflict level (H1; Fig. 2a). In LCE, approach RTs showed a clear graded pattern: decisions were slowest for HCT and fastest for NCT (Fig. 2b; HCT = 1.24 ± 0.02s, LCT = 1.20 ± 0.02s, NCT = 0.98 ± 0.01s). Pairwise t-tests confirmed significant differences between tree types (HC:NC t = 9.82, [0.20, 0.31], p < 0.001; LC:NC t = 10.1, [0.18, 0.26], p < 0.001), indicating that greater conflict increased decision difficulty.

**Figure 2:**
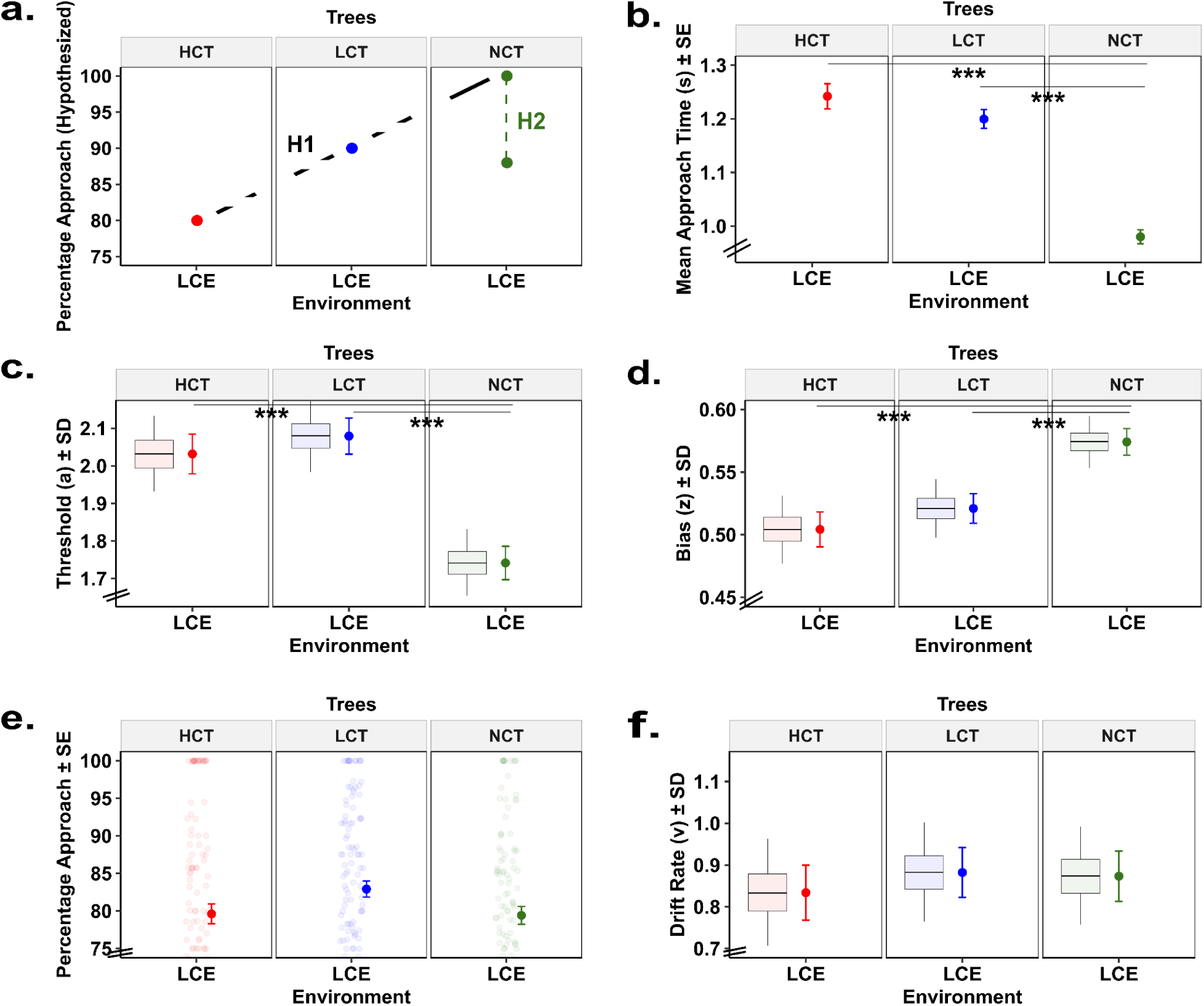
Hypothesized and observed effects of reward–punishment conflict in a low- conflict environment (LCE) a. Hypothesized mean percentage approach for high (red), low (blue) and no (green) conflict trees where hypothesis 1 shows sensitivity to local conflict whereas hypothesis 2 shows cross-conflict influence in LCE. b. Mean approach reaction times (in seconds ± SE, *standard error of mean*) for High, Low, and No conflict trees in LCE. c. Mean posterior estimates (± SD, *standard deviation*) of the threshold parameter ‘a’ from the Hierarchical Drift Diffusion Model (HDDM) for High, Low, and No conflict trees in LCE. d. Mean posterior estimates (± SD) of the starting bias parameter ‘z’ from the Hierarchical Drift Diffusion Model (HDDM) for High, Low, and No conflict trees in LCE. e. Mean percentage approach (± SEM) across LCE for High, Low, and No conflict trees. f. Mean posterior estimates (± SD) of the drift rate parameter ‘v’ from the Hierarchical Drift Diffusion Model (HDDM) for High, Low, and No conflict trees in LCE.

We fit a hierarchical drift–diffusion model (HDDM; see Methods) to examine the decision process. The response threshold *a*, indexing the caution exercised before committing to a choice, was significantly higher for HCT than NCT (HCT = 2.03 ± 0.05, LCT = 2.07 ± 0.04, NCT = 1.74 ± 0.04; Δmean = 0.287 [95% HDI: 0.158, 0.419], P(a(HCT.LCE) > a(NCT.LCE)) = 100.0%, q = 0.003; Fig. 2c). This confirms that high-conflict decisions are more difficult and require more deliberation.

The starting-point bias *z*, reflecting initial bias toward approach (*z* > 0.5) or avoidance (*z* < 0.5), also varied systematically with conflict. As expected, NCT elicited the strongest approach bias (z = 0.57 ± 0.01), which decreased for LCT (0.52 ± 0.01) and fell to neutrality for HCT (0.50 ± 0.01; Δmean = –0.284 [95% HDI: –0.430, –0.152], P(z(HCT.LCE) < z(NCT.LCE)) = 100.0%, q = 0.003; Fig. 2d). These patterns show that both decisional thresholds and initial biases track the level of conflict.

We next tested whether the approach to NCT would be near-universal (∼100%) if behaviour were governed solely by tree-specific contingencies (H1; Fig. 2a), or whether global environmental structure would suppress approach even in the absence of punishment (H2). The latter account was supported: approach rates were lowest for HCT (79.61 ± 1.32%) and slightly higher for LCT (82.91 ± 1.06%), but approach to NCT was unexpectedly reduced (79.42 ± 1.18%), matching HCT (Fig. 2e). This indicates a spillover effect in which conflict elsewhere in the environment discourages approach toward safe stimuli.

Finally, we examined whether the drift rate *v* captured the balance between instrumental value and Pavlovian avoidance. As predicted, drift rates were lowest for HCT (0.83 ± 0.06) and similarly low for NCT (0.87 ± 0.06), whereas LCT showed slightly higher values (0.88 ± 0.06; Fig. 2f). This pattern mirrors the behavioural avoidance observed for NCT and highlights the influence of global conflict structure on evidence accumulation.

Together, these findings demonstrate that approach–avoidance decisions are systematically shaped by reward–punishment conflict, reflected in RTs, decisional thresholds, and starting-point biases. Crucially, the reduced approach toward No Conflict trees captured both behaviourally and in drift dynamics indicates that choices are not determined by local contingencies alone but are tuned to broader environmental context. If avoidance in No Conflict trees stems from contextual rather than local factors, then altering global conflict should reshape approach behaviour. We tested this prediction in a second task block by manipulating environmental conflict.

### Environmental conflict modulates Pavlovian bias through instrumental control

We next asked whether the unexpected avoidance observed for No Conflict trees could be explained by the broader environmental structure. Specifically, we tested whether increasing the proportion of high conflict trees i.e., the level of *environmental conflict* would alter the expression of Pavlovian bias and shift behaviour toward instrumental responding. Under a High Conflict Environment (HCE), participants might show elevated avoidance overall; alternatively, heightened tonic arousal in such environments could suppress Pavlovian avoidance, thereby facilitating more goal-directed approach behaviour.

Behavioural data revealed systematic changes in approach behaviour across environments. In the HCE, approach rates decreased slightly for Low Conflict trees (81.45 ± 1.22%), increased for High Conflict trees (80.23 ± 1.32%), and increased markedly for No Conflict trees (82.52 ± 1.46%; Fig. 3a). A Wilcoxon rank-sum test confirmed a significant increase in approach for NCT (p = 0.019), and a Kruskal–Wallis test showed a trend-level effect of tree type (χ²(2) = 8.94, p = 0.10). This convergence across tree types suggests that global conflict attenuates Pavlovian bias, making choices less dependent on stimulus-specific reward–punishment contingencies.

**Figure 3:**
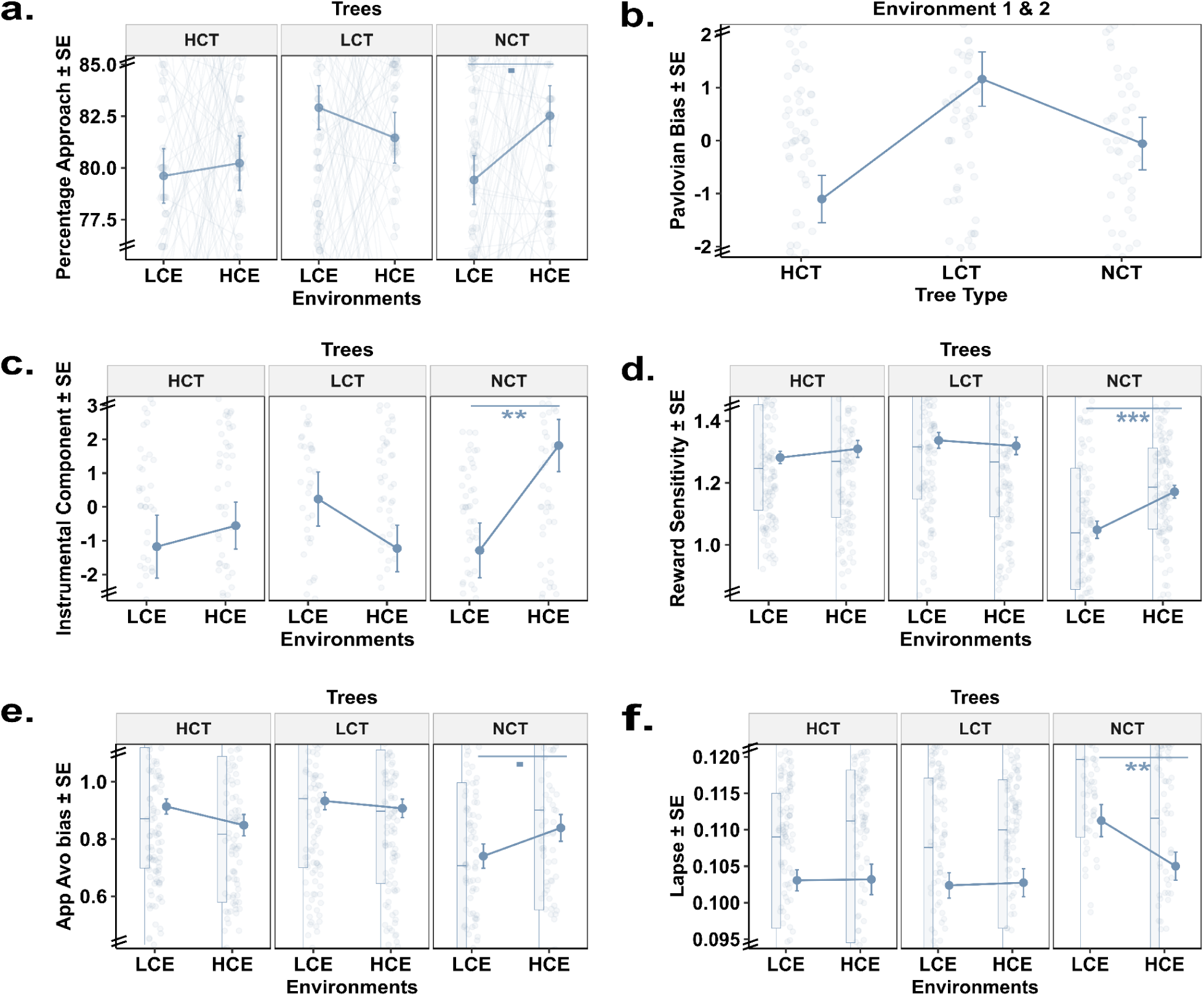
Behavioural and model-based measures across low and high conflict environments. a. Mean percentage approach (± SE) across LCE and HCE for High, Low, and No conflict trees. b. Pavlovian bias with SE, for High, Low and No conflict trees for Environment 1-2. c. Instrumental bias with SE, for High, Low and No conflict trees for LCE and HCE. d. Reward sensitivity parameter with SE from reinforcement learning model in LCE and HCE for High, Low and No conflict trees. e. Approach avoid bias parameter with SE from reinforcement learning model in LCE and HCE for High, Low and No conflict trees. f. Lapse parameter with SE from reinforcement learning model in LCE and HCE for High, Low and No conflict trees.

To decompose this effect, we separated approach behaviour into (i) a general Go bias, (ii) a Pavlovian component driven by specific tree type, and (iii) an Instrumental component reflecting residual, goal-directed variance (see Methods). Over and above a general Go bias and the tree-specific Pavlovian component (see *supplementary results*), we report that the residual Instrumental component for NCT increased significantly from LCE to HCE (–1.29 ± 0.81 → 1.81 ± 0.77; W = 5400.5, p = 0.0094; Fig. 3c), consistent with greater reliance on goal-directed action. Both behavioural percentages and model-based bias decomposition converge to show that environmental conflict suppresses Pavlovian influence in favour of instrumental control.

Computational modelling further supported this account. A 7-parameter reinforcement learning (RL) model captured the contributions of reward history, punishment history, Pavlovian bias, and choice noise (see Methods). Reward sensitivity reflecting the motivational impact of rewards was lowest for NCT during LCE (NCT: 1.05 ± 0.03; LCT: 1.34 ± 0.03; HCT: 1.28 ± 0.02). Post hoc Wilcoxon tests confirmed significant differences between LCT vs NCT (W = 10178, p < 9×10⁻¹¹) and NCT vs HCT (W = 3575, p < 3.3×10⁻⁹). In the HCE, reward sensitivity for NCT increased significantly (1.17 ± 0.02; W = 8587, p = 0.0004; Fig. 3d), consistent with re-engagement of reward motivation when Pavlovian avoidance had previously dominated.

The approach–avoidance bias parameter, here a reflection of the approach bias and therefore lower approach indicates higher avoidance bias, provides a direct computational index of Pavlovian influence. This parameter was non-zero for NCT and other trees in LCE supporting the existence of Pavlovian avoidance (0.74 ± 0.04 vs. LCT: 0.93 ± 0.03; HCT: 0.91 ± 0.03). Under HCE, this bias increased for NCT, meaning decrease in avoidance (0.84 ± 0.05; W = 7797, p = 0.03; post hoc p = 0.12; Fig. 3e), indicating reduced maladaptive avoidance.

The lapse parameter indexes choice randomness and decision noise, capturing the extent to which actions deviate from value-based predictions and again therefore measures Pavlovian bias. As expected from above results, the lapse parameter decreased significantly in NCT under HCE (0.1113 ± 0.0022 → 0.1050 ± 0.0019; W = 5174, p = 0.005; Fig. 3f). Thus, global conflict increased choice consistency, further implying stronger goal-directed control in HCE. The other parameters namely Punishment sensitivity, Punishment learning rate, Reward learning rate and action bias did not show any significant difference for NCT across environments (*supplementary results*, Fig. S1c, e, g, i respectively).

We next used hierarchical drift–diffusion modelling (HDDM) to jointly capture choice and RT patterns. Drift rates (v) were stable for HCT and LCT but increased selectively for NCT in HCE (0.87 ± 0.06 → 1.02 ± 0.08; Δmean = –0.155 [95% HDI: –0.349, 0.043], P(v(NCT.LCE) < v(NCT.HCE)) = 94.5%, q = 0.2; Fig. 4c), indicating enhanced instrumental value under high conflict.

**Figure 4:**
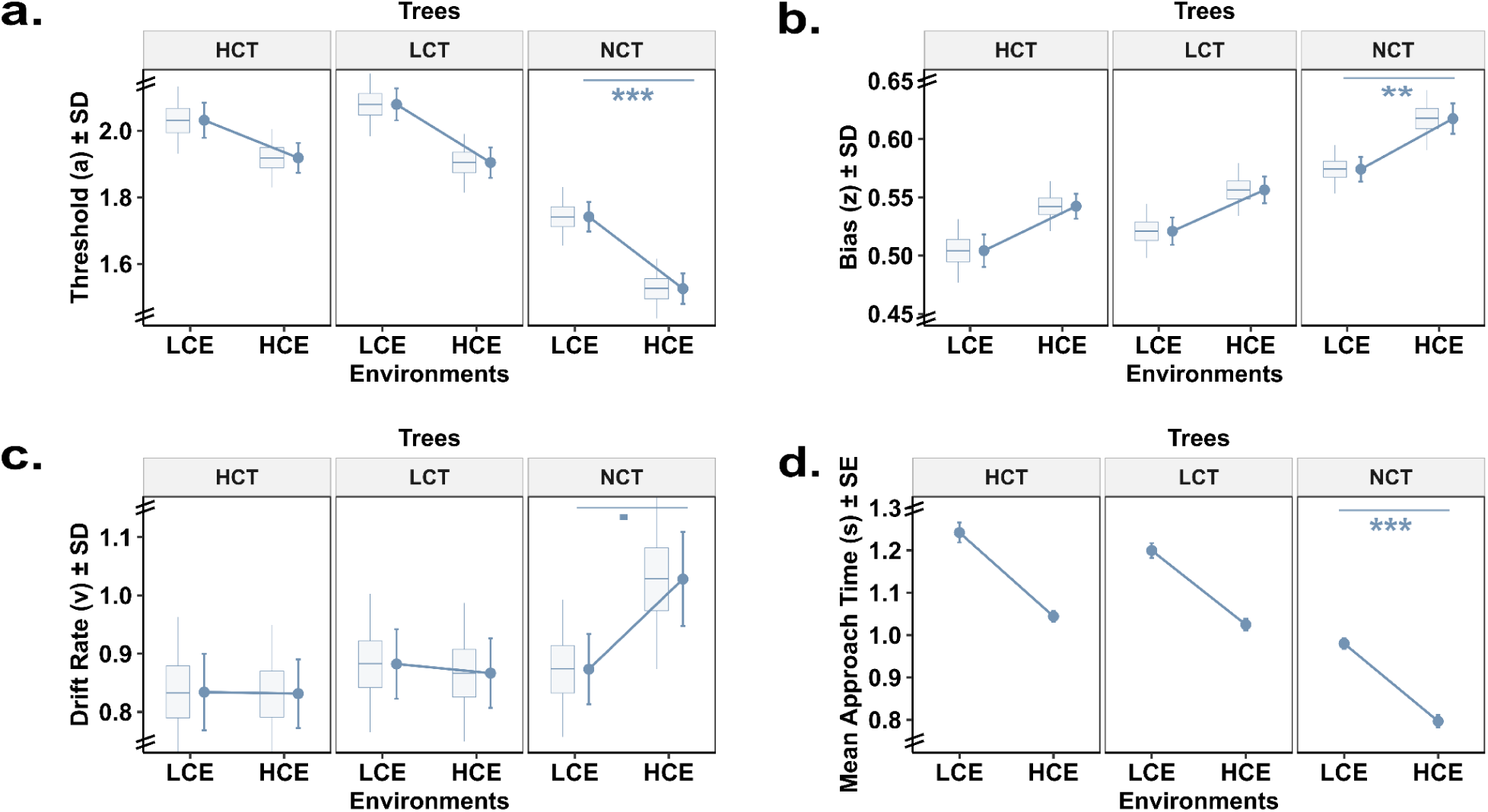
Hierarchical drift–diffusion model measures across low and high conflict environments. a. Mean posterior estimates (± SD) of the threshold parameter ‘a’ from the Hierarchical Drift Diffusion Model (HDDM) for High, Low, and No conflict stimuli in LCE and HCE. b. Mean posterior estimates (± SD) of the starting bias parameter ‘z’ from the Hierarchical Drift Diffusion Model (HDDM) for High, Low, and No conflict stimuli in LCE and HCE. c. Mean posterior estimates (± SD) of the drift rate parameter ‘v’ from the Hierarchical Drift Diffusion Model (HDDM) for High, Low, and No conflict stimuli in LCE and HCE. d. Mean approach reaction times (in seconds ± SE) for High, Low, and No conflict trees in LCE and HCE.

If environmental conflict truly modulated approach tendencies, then participants exposed to more High Conflict trees should exhibit reduced avoidance specifically for NCT. Because trials were pseudo-randomized, the proportion of HC encounters varied across individuals. This natural variability provided a test of whether environmental structure predicted changes in NCT avoidance. Indeed, increases in HC frequency significantly predicted greater reductions in NCT avoidance (β = –0.0053 ± 0.0017, t(114) = –3.14, p = 0.002). Thus, global conflict statistics, not just local trial contingencies, modulated avoidance expression.

Consistent with this, approach RTs, expected to slow under high conflict, were instead faster in the HCE (HCT: 1.04 ± 0.01s; LCT: 1.02 ± 0.01s; NCT: 0.80 ± 0.01s), with a significant decrease for NCT (t = 9.39, [0.14, 0.22], p < 0.001; Fig. 4d). Thresholds (a) decreased across tree types (HCT: 1.91 ± 0.04; LCT: 1.90 ± 0.01; NCT: 0.80 ± 0.01; Δmean = 0.215 [95% HDI: 0.089, 0.334], P(a(NCT.LCE) > a(NCT.HCE)) = 100.0%, q = 0.019; Fig. 4a). Starting-point biases (z) shifted toward approach in HCE, with the strongest shift for NCT (HCT: 0.54 ± 0.04; LCT: 0.55 ± 0.01; NCT: 0.61 ± 0.01; Δmean = –0.181 [95% HDI: –0.313, –0.042], P(z(NCT.LCE) < z(NCT.HCE)) = 99.4%, q = 0.06; Fig. 4b). Non-decision time(t) did not show any significant difference for NCT across environments (*supplementary results*, Fig. S1a).

Faster RTs, reduced thresholds, heightened reward sensitivity, and decreased choice noise all indicate greater arousal and attentional engagement under high conflict. Together, RL and HDDM analyses converge to show that global conflict reshapes the balance between Pavlovian and instrumental systems, reducing maladaptive avoidance and promoting more goal-directed behaviour precisely in contexts where local cues alone might otherwise bias decisions.

However, such benefits may not be uniform across individuals. Pavlovian avoidance varies substantially with trait anxiety, which biases individuals toward defensive responding even when rewards are available. This raises a key question: does environmental conflict confer the same instrumental override for high-anxious individuals, or does heightened avoidance blunt its modulatory effects? We turn to this next.

### Trait anxiety enhances Pavlovian avoidance but benefits disproportionately from environmental conflict

Trait anxiety is known to amplify avoidance in approach–avoidance conflict tasks. We therefore examined whether similar patterns emerged in our sample (N = 116; trait anxiety range: 26–72, median = 43). For illustration, we also divided participants into low (LTA; n = 64) and high trait anxiety (HTA; n = 52) groups, although all primary analyses treated trait anxiety dimensionally.

A linear regression confirmed that higher trait anxiety predicted reduced overall approach behaviour (β = –0.23, SE = 0.11, t(114) = –2.13, p = 0.035), consistent with stronger avoidance. In the Low Conflict Environment (LCE), this manifested most clearly for No Conflict (NCT) trees: HTA participants approached significantly less frequently than LTA (76.52 ± 1.88% vs. 81.77 ± 1.45%; t = 2.24, p = 0.026; Fig. 5a). Although this difference did not survive post hoc correction, it aligns with heightened Pavlovian avoidance in anxiety.

**Figure 5:**
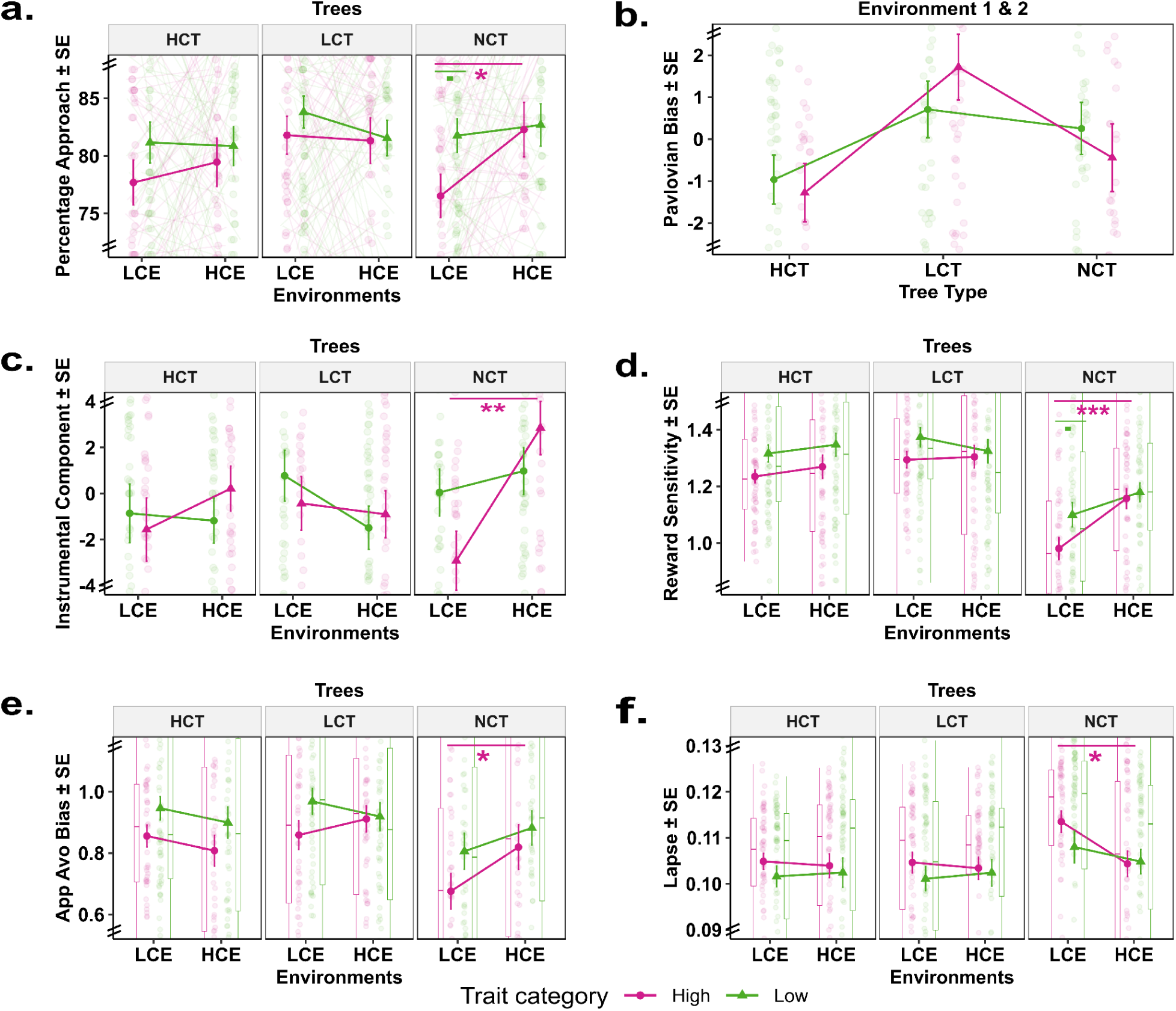
Behavioral and reinforcement-learning parameters across environments grouped by trait anxiety. a. Mean percentage approach (± SE) across LCE and HCE for High, Low, and No conflict trees, shown separately for high (pink) and low (green) trait anxiety groups. b. Pavlovian bias with SE, for High, Low and No conflict trees for Environment 1-2, shown separately for high (pink) and low (green) trait anxiety groups. c. Instrumental component with SE, for High, Low and No conflict trees for LCE and HCE, shown separately for high (pink) and low (green) trait anxiety groups. d. Reward sensitivity parameter with SE from reinforcement learning model in LCE and HCE for High, Low and No conflict trees, shown separately for high (pink) and low (green) trait anxiety groups. e. Approach-avoid bias parameter with SE from reinforcement learning model in LCE and HCE for High, Low and No conflict trees, shown separately for high (pink) and low (green) trait anxiety groups. f. Lapse parameter with SE from reinforcement learning model in LCE and HCE for High, Low and No conflict trees, shown separately for high (pink) and low (green) trait anxiety groups.

Crucially, this pattern reversed when environmental conflict increased. In the High Conflict Environment (HCE), HTA individuals showed a significant increase in NCT approach (76.52 ± 1.88% → 82.29 ± 2.35%; W = 1009.5, p = 0.026, r = 0.37), indicating reduced avoidance and enhanced goal-directed action (Fig. 5a). Moreover, over and above a general Go bias and Pavlovian component (see *Supplementary results*), the Instrumental component for NCT increased sharply in HTA from LCE to HCE (–2.92 ± 1.28 → 2.85 ± 1.16; W = 899, p = 0.003; Fig. 5c), reaching levels comparable to LTA. Thus, while anxiety heightened avoidance under low conflict, high environmental conflict appeared to normalize behaviour through stronger instrumental override.

Computational modeling using RL based models converged on the same conclusion. In LCE, reward sensitivity was lowest for NCT (1.05 ± 0.03) but increased significantly in HCE (1.17 ± 0.02; W = 850, p = 0.00046; Fig. 5d), particularly among HTA individuals. This reversal suggests that environmental conflict restored reward responsiveness precisely where avoidance was previously maladaptive.

Approach–avoidance bias, a direct RL measure of Pavlovian influence, was suppressed for NCT in LCE (0.74 ± 0.04 vs. HCT: 0.91 ± 0.03; LCT: 0.93 ± 0.03), but increased selectively for HTA in HCE (NCT: 0.84 ± 0.05; W = 1088, p = 0.046; Fig. 5e), signifying reduced avoidance. Additionally, lapse rates highest for HTA in LCE NCT (0.114 ± 0.00 vs. HCT: 0.105 ± 0.00; p = 0.005) declined in HCE (0.104 ± 0.00; W = 1761, p = 0.024; Fig. 5f), indicating reduced Pavlovian bias under high conflict. Punishment sensitivity, Punishment learning rate, Reward learning rate and action bias did not show any significant difference for NCT for the two groups: LTA and HTA across environments (*supplementary results*, Fig. S1d, f, h, j respectively).

HDDM parameters reinforced this pattern. Drift rates (v), indexing instrumental value, were lower in HTA than LTA in LCE for both HCT (0.75 ± 0.10 vs. 0.90 ± 0.08) and NCT (0.77 ± 0.08 vs. 0.96 ± 0.08). In HCE, drift rates increased across all tree types for HTA (HCT: 0.80 ± 0.08; LCT: 0.91 ± 0.09; NCT: 1.03 ± 0.10; Fig. 6c; Δv = –0.261 [95% HDI: –0.531, 0.042], posterior = 96.5%), indicating stronger instrumental drive.

**Figure 6:**
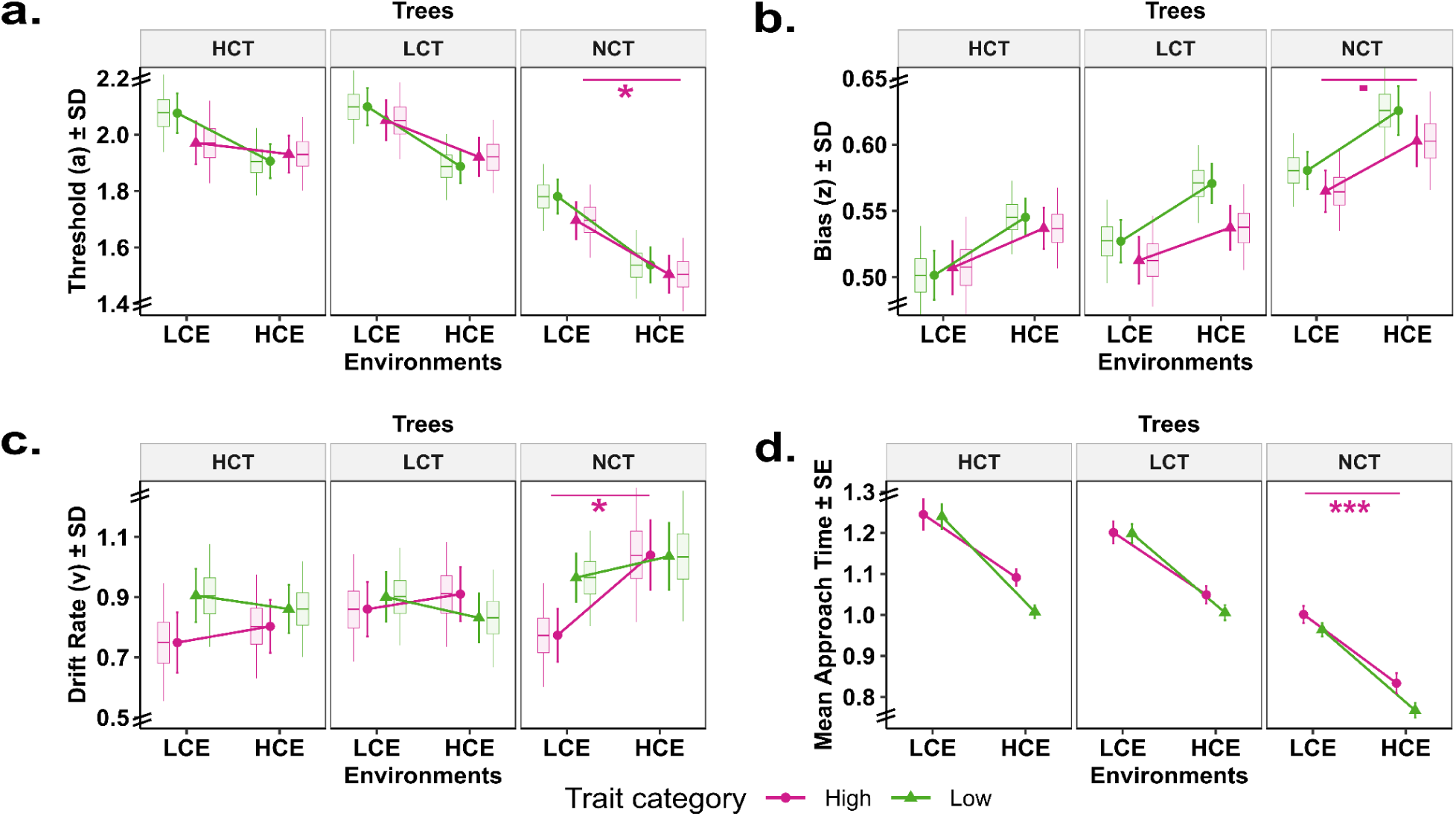
Drift–diffusion model measures across environments, grouped by trait anxiety. a. Mean posterior estimates (± SD) of the starting bias parameter ‘z’ from the Hierarchical Drift Diffusion Model (HDDM) for High, Low, and No conflict stimuli in LCE and HCE, shown separately for high (pink) and low (green) trait anxiety groups. b. Mean posterior estimates (± SD) of the threshold parameter ‘a’ from the Hierarchical Drift Diffusion Model (HDDM) for High, Low, and No conflict stimuli in LCE and HCE, shown separately for high (pink) and low (green) trait anxiety groups. c. Mean posterior estimates (± SD) of the drift rate parameter ‘v’ from the Hierarchical Drift Diffusion Model (HDDM) for High, Low, and No conflict stimuli in LCE and HCE, shown separately for high (pink) and low (green) trait anxiety groups. d. Mean approach reaction times (in seconds ± SE) for High, Low, and No conflict stimuli in LCE and HCE, shown separately for high (pink) and low (green) trait anxiety groups.

In LCE, HTA individuals exhibited elevated response thresholds (a) for conflict trees (HCT: 1.97 ± 0.07; LCT: 2.05 ± 0.07) relative to NCT (1.69 ± 0.06), reflecting overcautious evidence accumulation (Δa = 0.274 [95% HDI: 0.086, 0.457], posterior = 99.7%; Δa = 0.354 [95% HDI: 0.166, 0.532], posterior = 100.0%). In HCE, thresholds for NCT dropped markedly (1.50 ± 0.06; Fig. 6a; Δa = 0.189 [95% HDI: 0.025, 0.369], posterior = 98.7%), suggesting reduced indecision and faster commitment to approach behaviours.

Starting-point bias (z) showed a similar pattern. In LCE, HTA participants displayed minimal approach bias for conflict trials (HCT: 0.50 ± 0.02; LCT: 0.51 ± 0.01) but significantly greater bias for NCT (0.56 ± 0.01; Δz = –0.233 [95% HDI: –0.424, –0.026], posterior = 99.0%; Δz = –0.210 [95% HDI: –0.390, –0.018], posterior = 98.6%). Under HCE, approach bias increased further for NCT (0.60 ± 0.01; Fig. 6b), surpassing both conflict tree types (Δz = –0.161 [95% HDI: –0.365, 0.044], posterior = 94.5%). This suggests that global conflict amplified approach tendencies and counteracted avoidance in HTA individuals. Non-decision time(t) did not show any significant difference for NCT for the two groups across environments (*supplementary results*, Fig. S1b).

Approach latencies further supported this shift. RTs decreased significantly from LCE to HCE for NCT (HTA LCE: 1.25 ± 0.04; HTA HCE: 1.09 ± 0.02; LTA LCE: 1.24 ± 0.03; LTA HCE: 1.01 ± 0.02; Fig. 6d). HTA participants showed a clear speeding of NCT responses (median 0.76 s → 0.69 s; W = 739,453, p = 4.4×10⁻⁹). While LTA and HTA did not differ significantly in LCE (p > 0.10), they did in HCE (W = 451,827, p = 0.0038), indicating that high conflict selectively accelerated approach responses for HTA individuals.

Together, these findings reveal that trait anxiety magnifies Pavlovian avoidance under low-conflict conditions, yet high environmental conflict *disproportionately benefits* HTA individuals restoring reward sensitivity, reducing choice noise, enhancing approach motivation, and strengthening instrumental evidence accumulation. These results suggest that heightened environmental conflict, by elevating arousal and vigilance, can recalibrate maladaptive avoidance in anxiety and shift decision-making toward more adaptive, goal-directed strategies.

### Control experimental environments reveal boundary conditions of environmental influence

To test the generalizability of the environmental conflict effects, we examined behaviour in two additional control environments that manipulated cue–outcome associations and encounter rates.

In the first control environment, we maintained a high encounter rate of High Conflict (HC) trees but reversed the colour–tree mappings, thereby breaking previously learned cue–outcome associations while preserving global conflict statistics (HCE_R). Because the environmental structure remained unchanged, we predicted minimal behavioural disruption. As expected, choice patterns showed no re-emergence of Pavlovian bias, and both RL and HDDM parameters were statistically indistinguishable from those in the preceding HCE block (Figure S2). These findings indicate that under sustained high conflict, participants rely on stable instrumental policies that generalize even when stimulus–outcome mappings are reversed. Importantly, this stability was observed in both low- and high-anxiety groups.

In the next block, we reinstated a low-conflict environment (LCE_A) by reducing HC encounter frequency, as in the original LCE, while simultaneously increasing punishment probability (increased aversiveness). If punishment likelihood alone were sufficient to suppress Pavlovian bias, behaviour in LCE_A should have resembled the high-conflict environments (HCE and HCE_R). Instead, behaviour in LCE_A more closely tracked the reduced encounter rate: Pavlovian avoidance resurfaced, mirroring patterns from the initial LCE. Thus, encounter frequency, not punishment probability, was the primary driver of changes in Pavlovian bias. This pattern was largely consistent across anxiety groups, though HTA participants displayed slightly greater avoidance for both HC and NCT trees (Figure S3), aligning with generalized avoidance tendencies in anxiety.

Together, these analyses identify two key boundary conditions. When high conflict encounters remained frequent, performance was robust to cue–outcome reversal, reflecting dominance of instrumental control. When HC encounters were reduced, Pavlovian avoidance re-emerged, demonstrating sensitivity to global encounter statistics rather than to punishment magnitude per se.

### Environmental Conflict Affects Pavlovian Bias even in a Go-NoGo task

Lastly, to assess whether these effects extend beyond our task structure, we adapted a classical orthogonalized Go/NoGo task to incorporate low and high conflict environments (see *Supplementary Methods*). In the LCE, accuracy was significantly higher for congruent than incongruent trials (Median = 75.00% vs. 56.25%; Wilcoxon W = 3630.5, p < 0.001), demonstrating the expected Pavlovian bias on incongruent trials.

However, accuracy on incongruent trials improved significantly from LCE (M = 61.70%, SE = 3.02) to HCE (M = 70.63%, SE = 3.11; Figure S4a). A Wilcoxon signed-rank test confirmed a reliable environment effect (W = 2147.0, p = 0.023), with median accuracy increasing from 56.25% (LCE) to 73.04% (HCE). Although congruent accuracy remained descriptively higher, the advantage was no longer statistically significant, consistent with reduced Pavlovian interference in the HCE.

Trait anxiety did not significantly differentiate performance within either environment. Both LTA and HTA groups showed trend-level improvement for incongruent trials from LCE to HCE (LTA: 55.0 → 68.57, W = 468, p = 0.09; HTA: 57.5 → 80.3, W = 610.5, p = 0.1) and a trend-level group difference for congruent accuracy in HCE (LTA: 83.3, HTA: 90.0; W = 562.5, p = 0.1; Figure S4b).

### Deviations from Preregistered Predictions

Our preregistration outlined predictions that high trait anxious (HTA) individuals would show greater avoidance, impaired reversal learning, and reduced cognitive control. While HTA participants did exhibit overall elevated avoidance, the strongest effects emerged unexpectedly in No Conflict choices during low-conflict environments, rather than in high-conflict trials as preregistered. Under high conflict, avoidance differences between HTA and low trait anxious (LTA) individuals disappeared, indicating a dominant influence of global conflict structure. Contrary to predictions, reversal learning was unaffected by trait anxiety: cue–outcome reversal produced no group differences in behaviour or learning rates. Cognitive-control predictions were only partially supported: high-conflict environments improved behavioural alignment between HTA and LTA, whereas highly aversive contexts with reduced environmental conflict reinstated HTA avoidance, implying context-dependent control limitations. Overall, the main preregistered expectations were only partly met, HTA individuals showed elevated avoidance but not selectively under high conflict, nor impaired reversal learning, while exploratory analyses highlighted that Pavlovian biases, stronger in HTA individuals, were selectively suppressed under high conflict. These deviations underscore the central role of environmental structure in modulating anxiety-related choice biases.

## Discussion

Decision-making under uncertainty often reflects biases shaped by reward–punishment contingencies, yet less is known about how such biases are dynamically regulated (Bublatzky et al., 2017). Using a novel sequential approach–avoidance paradigm, we examined how environmental conflict influences Pavlovian biases and their modulation in individuals with varying trait anxiety.

Our findings show that approach behavior is sensitive to both immediate reward–punishment conflict and the broader environmental context. At the behavioral level, higher punishment increased avoidance, reflected in drift diffusion model (DDM) parameters, higher thresholds (a) and reduced approach bias (z) indicating greater decision difficulty and diminished approach. Critically, however, the broader environmental conflict exerted an additional influence: exposure to conflict trees suppressed approach even toward No Conflict trees, consistent with Pavlovian bias. This effect was exaggerated in individuals with higher trait anxiety, aligning with prior evidence linking anxiety to heightened avoidance (Qi et al., 2018; Hofmann & Hay, 2018; Ball & Gunaydin, 2022).

Importantly, Pavlovian bias was not static. Increasing environmental conflict attenuated its influence, accompanied by enhanced value sensitivity (higher drift rates and reward sensitivity) and reduced lapse rates. This suggests that arousal- and attention-related mechanisms may boost the salience of instrumental value representations, thereby supporting goal-directed responding.

An alternative explanation is that participants strategically favored high-conflict trees, assuming these would yield more valuable outcomes in the high conflict environment. Yet this fails to explain the data: avoidance of No Conflict trees should have persisted or increased across environments, but approach to these trees rose in high conflict environment. A generalized action/ Go bias offers another possibility, supported by faster responses, lower thresholds, and higher starting point bias in DDM parameters (Pedersen et al., 2021). However, such a global tendency would predict uniform increases in approach, whereas our effects were tree-specific, linked to drift rate and lapse parameters. Incomplete learning of contingencies is also unlikely. Pavlovian bias re-emerged in block 4 when environmental conflict reduced again. Taken together, while these alternatives provide partial explanations, the most parsimonious account is that Pavlovian bias, modulated by environmental conflict, best captures the observed dynamics.

Prior work has consistently demonstrated the influence of Pavlovian biases but rarely their modulation. Only a handful of studies have reported reductions in bias, either through extended training or semantic contexts (Ereira et al., 2021; Fleming et al., 2023). Our paradigm advances this literature by showing that global conflict can suppress Pavlovian influence in active avoidance settings.

A key question is whether the reduction in Pavlovian bias we observed is unique to our sequential tree paradigm or whether it generalizes to more established approaches. To test this, we implemented an orthogonalized Go/NoGo design, a widely used paradigm for probing conflicts between Pavlovian and instrumental control. In these tasks, Pavlovian reward-seeking can conflict with instructed NoGo responses, while punishment avoidance can conflict with required Go responses (Guitart-Masip et al., 2012). Consistent with our hypothesis, high-conflict contexts significantly improved accuracy for Pavlovian-incongruent cues, mirroring the pattern seen in our sequential task. This replication is important, as prior Go/NoGo studies have largely emphasized the robustness of Pavlovian biases but have shown limited evidence for their contextual modulation (Finotti et al., 2025; Geurts et al., 2013; Huys et al., 2011; Saeedpour et al., 2023). By demonstrating that environmental conflict can attenuate bias across both novel and classical tasks, our findings extend beyond earlier work and provide convergent evidence for a broader, generalizable mechanism through which context regulates the expression of Pavlovian influence.

The observed dynamics can be interpreted within the adaptive gain theory of the locus coeruleus–norepinephrine (LC–NE) system (Aston-Jones & Cohen, 2005). This framework posits that tonic and phasic LC activity regulate the balance between exploration and exploitation, thereby shaping decision-making and cognitive flexibility (Jepma, 2010; Maness et al., 2022). We propose that environmental conflict modulates tonic LC activity: in low-conflict environments, suboptimal NE tone leaves instrumental systems under-engaged, allowing Pavlovian biases to dominate. In contrast, high-conflict contexts may elevate tonic NE toward an optimal range, improving the signal-to-noise ratio of value representations and supporting goal-directed control (Fig 7a). This interpretation is consistent with our behavioral findings, where high conflict was associated with increased drift rates and reduced lapses, reflecting enhanced value sensitivity and greater reliance on instrumental processes.

**Figure 7:**
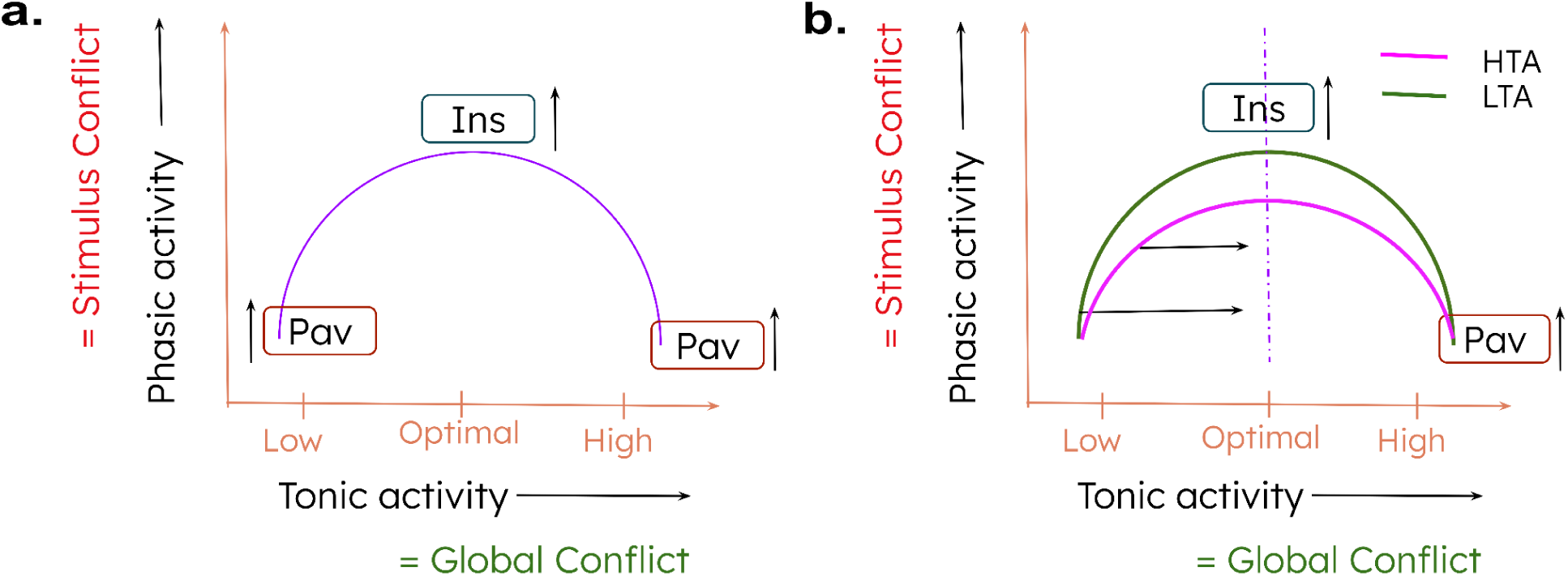
Schematic model of arousal-dependent modulation of Pavlovian bias. a. The Yerkes-Dodson model of arousal and performance under LC-NE framework to explain tuning of Pavlovian to Instrumental responding whereby baseline tonic activity of LC neurons sets the stage for baseline response which is influenced by the global level of conflict whereas phasic activity of LC neurons is controlled by stimulus-specific conflict. A combination of these two responses determine Pavlovian-to-Instrumental switching. Low and high tonic activity despite high phasic activity fails to elicit optimal response thus favoring Pavlovian response. However, optimal tonic activity with appropriate stimulus conflict can lead to optimal instrumental response. b. The above model also accounts for high (pink) and low (green) trait anxiety group behavior, whereby high trait anxiety has a shallower response curve leading to higher susceptibility to tonic activity and in turn global level conflict.

Group-level differences strengthen this account. High trait anxious individuals, exhibited exaggerated Pavlovian bias under low-conflict conditions (Mkrtchian et al., 2017). Yet, they benefited most from high-conflict environments, where increased arousal likely suppressed maladaptive biases and redirected attention toward task-relevant value signals. This pattern aligns with the Yerkes–Dodson principle: performance improves at intermediate arousal levels but deteriorates when arousal is either too low or too high (Fig 7b). For anxious individuals, whose baseline arousal states may be suboptimal, environmental conflict may push tonic NE activity into a more advantageous range, thereby enabling more effective instrumental control.

Looking ahead, future studies incorporating physiological markers such as pupillometry or direct indices of norepinephrinergic function will be critical to testing the proposed LC–NE mechanisms more directly. By situating these effects within a neurobiological framework of arousal and control, our results demonstrate that environmental conflict dynamically regulates Pavlovian bias in approach–avoidance decisions. This has particularly strong implications for understanding anxiety, where maladaptive avoidance behaviors may be driven by overreliance on Pavlovian systems. These findings suggest translational pathways: carefully modulated conflict exposure, cognitive training under controlled arousal states, or neuromodulatory approaches targeting LC–NE tone (e.g., tACS, pharmacological agents) may help rebalance the interplay between Pavlovian and instrumental systems. Such strategies could provide novel means of reducing maladaptive avoidance and enhancing adaptive decision-making in anxiety.

## Acknowledgment

This project was supported by grants from the DBT-Wellcome Trust India Alliance and the IIT Kanpur Startup Grant awarded to Arjun Ramakrishnan. We gratefully acknowledge the Council of Scientific and Industrial Research (CSIR) for awarding a Senior Research Fellowship (SRF) to Priyanshu. We also thank Advaith Kannan for developing the initial version of the game in PsychoPy, Kritnandan for streamlining the reinforcement learning model implementation in Stan, and Shalem Rajkumar for contributing to the initial data structure pipeline during his summer internship.

## Competing interests

The authors declare no competing interests.

## Author contributions

Priyanshu conceptualized the study, designed the task, collected and analyzed the data, and wrote the initial draft of the manuscript. Arjun Ramakrishnan supervised the project, contributed to the conceptual framework, provided critical revisions, and guided the interpretation of results. Both authors discussed the findings, edited the manuscript, and approved the final version.

## Methods

### Ethics statement

The Institutional Ethics Committee (IEC), Indian Institute of Technology Kanpur (IITK) (IITK/IEC/2024-25/II/36) approved all procedures and performed protocol in accordance with the obtained approval. Participants e-signed informed consent before participating.

### Power-Analysis for sample size

We performed power analysis for generalized linear mixed effect regression (glmer) models using SIMR package in R (Green & MacLeod, 2016). Power analysis revealed 46% power for an effect size −0.026 in 24 participants. We extrapolated the results for effect size −0.017 in 70 participants and got 62% power. Based on these results, we decided to collect data from further 70 participants, totalling up to ∼140 participants which should be a good number to get power above 80% for that effect size.

### Participants

Participants were recruited through institute wide email from Indian Institute of Technology Kanpur and consisted mostly of the student population of IIT Kanpur. Total 140 participants participated in the experiment (**aged - mean: 22.48, sd: 5.32**); (**Females: 39**) and were assigned to either variant of the game i.e Red or Green variant. Of which 24 participants were excluded based on 3 exclusion criterias: Technical or game related crashes or didn’t understand game instructions and lastly outliers based on LOOIC, final count was 116 participants (**aged - mean: 22, sd: 4.5**); (**Females: 32**). Participants received financial compensation based on their performance upon completion of the experiment.

### Approach-avoidance conflict paradigm

#### Design

A novel sequential approach-avoidance task designed to understand how reward-punishment conflict and environment affect approach avoidance decisions. Task consists of three colored trees, each with different magnitudes of reward (R) and punishment (P) thus varying degrees of conflict: 1R/0P (No conflict tree), 2R/1P (Low conflict tree) and 3R/2P (High conflict tree) (Figure 1b). To add to the complexities, we have four environments referred to as Blocks which differ in availability of No conflict/ High conflict trees and aversiveness (Table 1, Figure 1c). The frequency of HC trees in a block determines conflict at the level of environment referred to as Environmental conflict. For Ex:Block I which has higher frequency of NC and lower frequency of HC is called “Low Conflict Environment” whereas HCE is a “High Conflict Environment”. The Block transition from Block II to III includes reversal of contingencies between tree color and conflict i.e. Red tree becomes NC and Green tree becomes HC (High Conflict Environment_Reversal, HCE_R), whereas in Block IV we increase aversiveness i.e. punishment encounter rate from 0.3 to 0.7 in conflict trees (Low Conflict Environment_Aversive, LCE_A). Each block lasted 5 minutes while the total duration of the game was 20 minutes.

**Table 1:** Block-wise breakup of frequency and reward-punishment contingencies in each tree type.

| **Environment/Trees \|\|**<br>**Aversiveness** | **No Conflict Tree**<br>**(1R/0P)** | **Low Conflict Tree**<br>**(2R/1P)** | **High Conflict Tree**<br>**(3R/2P)** |
| --- | --- | --- | --- |
| **Block I (LCE) \|\| 0.3** | 0.45 | 0.35 | 0.20 |
| **Block II (HCE) \|\| 0.3** | 0.20 | 0.35 | 0.45 |
| **Block III (HCE_R) \|\| 0.3** | 0.20 | 0.35 | 0.45 |
| **Block IV (LCE_A) \|\| 0.7** | 0.45 | 0.35 | 0.20 |
| **Reward/Punishment** | 1-10/None | 11-20/-1-(-10) + S | 21-30/-11-(-20) + SS |

### Procedure

#### a. Baseline testing for Shock

A shock work-up procedure was performed to adjust shock intensities individually for each participant based on their shock sensitivity/tolerance. A single shock of pulse width 200 *μ*s of different intensity was delivered to non-dominant wrist via a pair of silver chloride electrodes using a DS8R stimulator (Digitimer Ltd, Welwyn Garden City, UK). Participants received shock sequentially with a 0.5 step increase in amplitude (starting with 0mA and moving up), which they had to rate on a scale from 1 to 5 ( 1 meaning “barely felt” and 5 “shocking intensity turning painful”). Participant’s rating of 1 (barely felt), 3 (very slightly irritating), 4 (highly irritating) and 5 (painful) were recorded separately to be used in the game.

#### **b.** Pre-game survey

Participants’ details were recorded through an online Qualtrics platform along with reading & e-signing of consent form, followed by some standard self-reported questionnaires like Spielberger’s State Anxiety Inventory (STAI-Y1) and Spielberger’s Trait Anxiety Inventory (STAI-Y2) (Spielberger, 1983), Generalized Anxiety Disorder 7 (GAD7), Patient Health Questionnaire 9 (PHQ9), Intolerance of Uncertainty (IUS-12) and Barratt Impulsiveness Scale - 11.

Trait anxiety was assessed using questionnaire scores, which were either analyzed continuously in relation to task measures or used to divide participants into high and low anxiety groups. In the present paper, we focus on trait anxiety scores as the primary measure. Group scores ranged from 26 to 72 (Mean: 42.9, SD: 8.7), with a median of 43 (Figure S5).

#### **c.** Game play

The game was designed and run on Psychopy (Version v2022.2.5). Participants performed a simple two alternative choice task when they were first presented with a neutral white tree and they had to click on the reveal button to reveal one of the three types of stimuli represented by green, blue and red coloured trees. Simultaneously, with the colored trees, two options were presented: “Harvest” and “Skip”. Participants could choose to approach (harvest) or avoid (skip) at each trial for each tree. On harvesting they either get a reward points (fruits) or punishment as points loss and shock (thorns) while on skipping they move on to the next trial (Figure 1b). The objective for participants in the game was to maximize reward points(fruit collection) to get maximum monetary bonus reward. The game lasted for 20 mins.

#### **d.** Post game survey

We collected post game feedback about the experiment, which included questions that checked for their attentiveness and learning and motivation for playing the game.

### Pre-registration

This study was pre-registered on the AsPredicted platform (Wharton Credibility Lab) (ID: *AsPredicted#153022*) prior to data collection. The pre-registration document includes the study hypotheses, experimental design, sample size determination, stimuli details (red, blue, and green trees for High, Low, and No conflict stimuli and counterbalancing), reward and punishment structure, and planned statistical analyses, including the use of a percentage approach to measure behavior and comparisons between high and low trait anxiety groups.

The pre-registration can be accessed at [https://aspredicted.org/8xcq-nhv8.pdf]. Any deviation, whatsoever, is in expected results and their interpretation, the methods remain the same.

### Model Agnostic Behavioral Analysis

All major data analysis was done on R (Version: 4.3.1) and RStudio (Version: 2024.12.0+467). We did Pearson’s correlation test, t-test and ANOVA using R packages.

We assessed the distribution of all variables of interest (e.g., percentage approach, instrumental bias) across blocks and conflict levels. Normality was evaluated using three approaches: visual inspection of histograms, Q–Q plots of residuals, and the Shapiro–Wilk test. Homogeneity of variance was assessed using Levene’s test. If both assumptions of normality and homogeneity were met, we performed parametric tests, including pairwise *t*-tests and ANOVA with Tukey HSD post hoc comparisons. When assumptions were violated, we first applied bootstrapping procedures. If bootstrapping did not yield reliable results, we conducted non-parametric tests, specifically the Wilcoxon rank-sum test and the Kruskal–Wallis test, followed by Benjamini–Hochberg (BH) corrected post hoc comparisons.

We specified linear, logistic and beta regression models as well as mixed effect models by predicting decisions as a function of Trees, Blocks and Trait anxiety scores using lme4 and betareg package, utilizing lm(), glmer() and betareg() in R.

For example:

Model_1 = lm(Trait Score ∼ Percentage approach)

Model_2 = betareg( *Δ NC avoidance proportion (T2-T1) ∼ Δ HC Frequency (T2-T1) )*

Model_3= glmer(Decision ∼ Conflict + Block + (1|Participant), family = “binomial”)

Linear regression(lm) measures the relationship between a continuous dependent variable and one or more independent variables by fitting a straight-line equation using ordinary least squares (OLS).

Logistic mixed-effects models (glmer) estimate the probability of a binary outcome (0/1) while accounting for both fixed (predictors) and random effects (group-level variations), using maximum likelihood estimation (MLE) to model log-odds of the outcome and incorporates random intercepts/slopes for hierarchical data.

Beta regression (betareg) is designed for modeling proportions or rates (value between 0 and 1) by assuming a Beta distribution and using MLE to estimate both the mean and precision of the response variable.

We checked for model assumptions for each model type, for example: normality of residuals using QQ plots and the Shapiro-Wilk test, homoscedasticity was checked with residuals vs. fitted plots and multicollinearity was examined using Variance Inflation Factor (VIF). For beta regression (betareg), heteroskedasticity and non-linearity were examined using diagnostic residual plots, and the link function choice was validated using the Akaike Information Criterion (AIC). These checks ensured that model assumptions were met, improving the reliability of the analyses.

### Approach-avoidance measure

We quantified approach-avoidance decisions in the task as *percentage approach metric* across different tree types (Conflict levels: No Conflict, Low Conflict, High Conflict) and blocks (Environments) with the data collected from 116 participants:

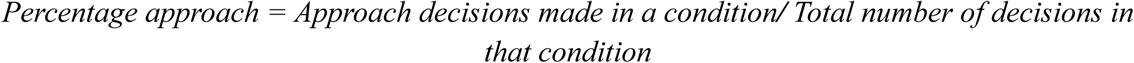

The reward associated with different tree (conflict) type is as follows: No Conflict: Small Reward , No Punishment (1R/0P)

Low Conflict: Double Reward, Low Punishment (2R/1P) + 3-point rated shock

High Conflict: Triple Reward, High Punishment (3R/2P) + 4-point rated shock

### Bias construction

**Go bias:** This relates to a general motivation to act towards approach options. There is one approach bias for each group, LTA and HTA. It was calculated using following formula:

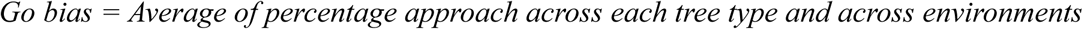

#### Pavlovian component

This relates to tendency to choose either approach or avoid based on stimulus i.e. tree color or conflict type. Thus, there are three Pavlovian biases for each group: P_NC, P_LC and P_HC. It was calculated using following formula:

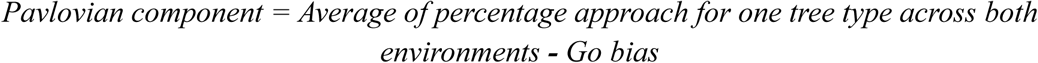

The positive Pavlovian component represents Pavlovian approach bias whereas negative Pavlovian component represents Pavlovian avoidance bias.

#### Instrumental component

This relates to tendency to choose either approach or avoid based on reward-punishment learning for each tree type in four blocks. Thus, there are total 12 Instrumental biases for each group: I_T1_NC, I_T1_LC, I_T1_HC, I_T2_NC, I_T2_LC, I_T2_HC, I_T3_NC, I_T3_LC, I_T3_HC, I_T4_NC, I_T4_LC and I_T1_HC. It was calculated using following formula:

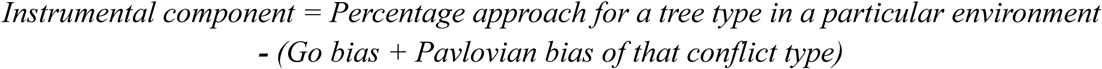

Positive Instrumental component represents instrumentally learnt actions whereas negative bias represents Pavlovian influence over actions.

Here we assume that the Pavlovian component across a particular tree type remains the same across two environments and therefore the decisions to approach or avoid are influenced by Instrumental learning of reward and punishment outcomes.

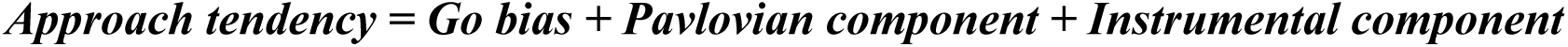

### Drift Diffusion Modelling

To gain insight into the dynamics of decision-making, we employed the Hierarchical Drift Diffusion model (HDDM 0.8.0) (Wiecki et al., 2013), a package in python3 using reaction time and choice data. Model fitting was performed using Markov chain Monte Carlo (MCMC) sampling with 5,000 samples and a burn-in of 500 samples.

The drift diffusion model is a model of the cognitive processes involved in simple two-choice decisions. The diffusion model assumes that decisions are made by a noisy process that accumulates information over time from the starting point towards one of the two response boundaries in our case approach and avoid. It falls in the category of models that take speed-accuracy trade off into account, called Sequential Sampling model (SSM), conceptualizing decision making as a process of noisy evidence accumulation (Forstmann et al., 2016; Ratcliff et al., 2016). A response is generated when the accumulated evidence reaches a decision boundary, and the time for accumulating the decision evidence is the reaction time (RT). DDM decomposes joint distribution of RT and choice data into several cognitively meaningful parameters i.e. drift rate (v) for the speed of accumulating evidence, decision boundary/ threshold (a) for decision caution, the starting point (z) for evidence accumulation bias, and non-decision time for motor response (t). The starting point is labelled as the “z” parameter. The z value of 0.5 reflects no starting point bias towards either of the boundaries. The distance between response boundaries are labelled as “a” and bounded from 0 to a. The rate of accumulation of information is called the drift rate “v” and is determined by the quality of stimulus information or certainty of a particular decision.

We fitted several HDDM models on our data. Here, we report two best models based on DIC score comparison (see *supplementary methods*), one with Trait category dependency and another without to see group level differences between predictors with and with trait categorization allowing all four model parameters: a, t, v and z to vary by these specified groups.

**hddm_model_1** = hddm.HDDM(data, bias = True, depends_on={’a’: ‘Conflict’:’Block’, ‘v’: ‘Conflict’:’Block’, ‘z’: ‘Conflict’:’Block’, ‘t’: ‘Conflict’:’Block’}

**hddm_model_2** = hddm.HDDM(data, bias = True, depends_on={’a’: ‘Conflict’:’Block’:’Trait_category’, ‘v’: ‘Conflict’:’Block’:’Trait_category’, ‘z’: ‘Conflict’:’Block’:’Trait_category’, ‘t’: ‘Conflict’:’Block’:’Trait_category’}

To ensure that the model had converged to a stable solution prior to interpreting its parameters, we computed the Gelman–Rubin convergence statistic (Gelman & Rubin, 1992). The maximum Ȓ value across runs was 1.01, indicating satisfactory convergence. The final model was estimated using 5000 posterior samples (with 500 burn-in iterations). Visual inspection of the posterior trace plots further confirmed convergence. Although several models were tested, the two reported here demonstrated the lowest DIC values.

Before implementing the HDDM models, we performed rigorous data filtering at the participant level to remove outliers. As specified in the preregistration, trials with response times exceeding the mean + 5 SD were excluded from further analyses.

### Approach-avoidance reinforcement learning model

Behaviour was modelled using a seven-parameter approach–avoidance reinforcement-learning model adapted from established Go/NoGo frameworks (Mkrtchian et al., 2017) modified to accommodate active approach and avoidance decisions. However, we first fitted a set of hierarchical Bayesian reinforcement-learning models using the hBayesDM package (Ahn et al., 2017) in R to characterize basic reward and punishment learning dynamics in the task (see *supplementary methods*).

The model separately estimated learning rates for reward and punishment outcomes (α_reward, α_punishment), outcome sensitivity parameters for reward and punishment, a general action bias, an approach–avoidance bias that scaled with state value, and a lapse parameter capturing random choice behaviour. Action values were updated using a sensitivity-scaled delta rule applied to both action-specific and state values. For approach (go) actions, the action weight combined the learned value with a general action bias and a Pavlovian approach–avoidance term proportional to state value, whereas avoidance (nogo) actions depended solely on their learned value. Choices were generated via a softmax function with a lapse mixture to account for stimulus-independent responding.

Learning rates and lapse parameters were constrained to the unit interval using beta priors, while sensitivity and bias parameters were assigned normal priors. Model fitting used a factorial prior structure (Block × Conflict × Trait category; 24 priors).

**Table 2:** Reinforcement-learning model parameters and priors.

| Parameters | Distribution | The range of Distribution value lies |
| --- | --- | --- |
| alpha_reward | beta(2, 18) | [0, 1] |
| alpha_punishment | beta(2, 18) | [0, 1] |
| sensitivity_reward | normal(1, 1) | $(-\infty, +\infty)$ |
| sensitivity_punishment | normal(1, 1) | $(-\infty, +\infty)$ |
| action_bias | normal(1, 1) | $(-\infty, +\infty)$ |
| app_avo_bias | normal(1, 1) | $(-\infty, +\infty)$ |
| lapse | beta(2, 18) | [0, 1] |

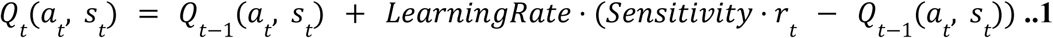

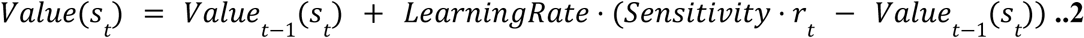

**For action a = harvest (go):**

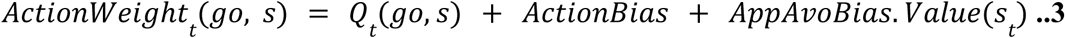

**For action a = skip (nogo):**

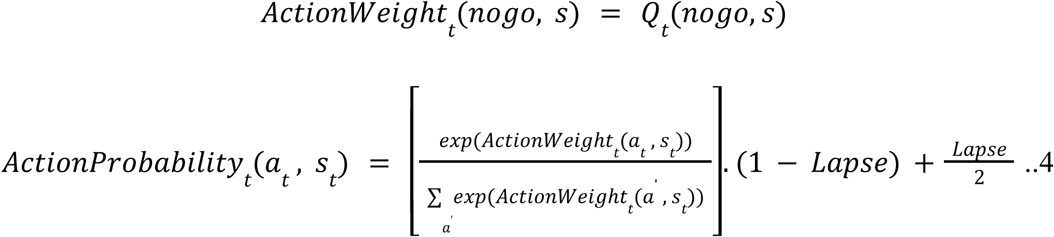

## Supplementary Material

Table of Content:

Supplementary Methods

i. HDDM model comparison
ii. hBayesdM: Hierarchical Bayesian model of Decision Making
iii. Control Experiment II

Supplementary Results

i. Bias construction (supplementary):

Supplementary Tables:

Table S1: DIC scores comparison across HDDM models

Table S2: hBayesDM Model Specifications

Table S3: LOOIC score comparison for different hBayesDM models

Table S4: LOOIC score comparison for different priors of winning hBayesDM model

Table S5: Block-design for Experiment II (Orthogonalized Go-NoGo task) paradigm Table S6: Task outcome contingencies

Supplementary Figures:

Figure S1: Additional hierarchical drift–diffusion and reinforcement-learning model parameters

Figure S2: Behavioural and computational signatures across cue–outcome reversal in high-conflict environment

Figure S3: Behavioural and computational patterns in a low-conflict but aversive environment

Figure S4: Effects of environmental conflict on Go/NoGo task performance

Figure S5: Distribution of trait anxiety scores with median-based grouping across experimental samples

## Notes

### Competing Interest Statement

The authors have declared no competing interest.

