## Supplementary Material for "Environmental Conflict Modulates Pavlovian Bias"

**Table of Content:**

A. Supplementary Methods
  i. HDDM model comparison
  ii. hBayesdM: Hierarchical Bayesian model of Decision Making
  iii. Control Experiment II

B. Supplementary Results
  i. Bias construction (supplementary):

***Supplementary Tables:***

Table S1: DIC scores comparison across HDDM models
Table S2: hBayesDM Model Specifications
Table S3: LOOIC score comparison for different hBayesDM models
Table S4: LOOIC score comparison for different priors of winning hBayesDM model
Table S5: Block-design for Experiment II (Orthogonalized Go-NoGo task) paradigm
Table S6: Task outcome contingencies

***Supplementary Figures:***

Figure S1: Additional hierarchical drift–diffusion and reinforcement-learning model parameters
Figure S2: Behavioural and computational signatures across cue–outcome reversal in high-conflict environment
Figure S3: Behavioural and computational patterns in a low-conflict but aversive environment
Figure S4: Effects of environmental conflict on Go/NoGo task performance
Figure S5: Distribution of trait anxiety scores with median-based grouping across experimental samples

### A. Methods:
#### i. HDDM Model comparison:

We compared a set of hierarchical drift–diffusion models that differed in whether decision parameters were modulated by environmental conflict, block structure, and trait anxiety (Table S1).

| Models | DIC Scores |
|---|---|
| 1. Only Trait category | 71146.660603 |
| 2. Only Conflict | 68499.042237 |
| 3. Only Block | 65819.020790 |
| **4. Conflict + Block** | **62264.741665** |
| 5. Block+ Trait category | 65816.029093 |
| **6. Conflict + Block + Trait category** | **62271.156316** |
| 7. Conflict* Trait category + Block | 68896.180914 |
| 8. Block*Trait category + Conflict | 68864.024985 |
| Lumped model DIC | 72311.983390 |

***Table S1:*** *DIC scores comparison across HDDM models, reported models have been highlighted in bold with lowest DIC scores.*

**ii. hBayesdM: Hierarchical Bayesian model of Decision Making :**
We first fitted a set of hierarchical Bayesian reinforcement-learning models implemented in the hBayesDM package (Ahn et al., 2017) in R to characterize basic reward and punishment learning dynamics in the task (Table S2). These models, which were estimated using Stan with four chains (2,000 burn-in samples and 4,000 retained samples per chain), capture learning rates, outcome sensitivity, and choice stochasticity, and were fitted to trial-wise choice, gain, and loss data for each participant.

Across models, parameter estimates revealed learning dynamics consistent with those obtained using our primary analyses. However, the hBayesDM model family does not include an explicit parameter indexing Pavlovian approach–avoidance bias, which was central to our theoretical questions regarding the modulation of approach–avoidance behaviour by environmental structure. We therefore extended our analyses by fitting a seven-parameter approach–avoidance reinforcement-learning model that explicitly dissociates instrumental learning from Pavlovian approach–avoidance influences.

| **Model** | **NP** | **Parameters** | | | | | |
|---|---|---|---|---|---|---|---|
| banditNarm_4par | 4 | Reward Sensitivity | Punishment Sensitivity | Reward Learning Rate | Punishment Learning Rate | | |
| banditNarm_lapse | 5 | Reward Sensitivity | Punishment Sensitivity | Reward Learning Rate | Punishment Learning Rate | Lapse | |
| banditNarm_lapse_decay | 6 | Reward Sensitivity | Punishment Sensitivity | Reward Learning Rate | Punishment Learning Rate | Lapse | Decay |
| banditNarm_singleA_lapse | 4 | Reward Sensitivity | Punishment Sensitivity | Learning Rate | | Lapse | |

***Table S2:*** *hBayesDM Model Specifications.*

*We fitted 4 models using the hbayesDM package. NP = number of parameters. Model = model names implemented in the hbayesDM package.*

The banditNarm models (where i refers to a given bandit, t refers to trial) were calculated by inputting reward and punishment values separately to the following equations:

$$Value^{rew}_{t(i)} = Value^{rew}_{t(i)} + LearningRate_{rew} * Prediction\ Error^{rew}_{t(i)} \quad ..1$$

$$Value^{pun}_{t(i)} = Value^{pun}_{t(i)} + LearningRate_{pun} * Prediction\ Error^{pun}_{t(i)} \quad ..2$$

$$Prediction\ Error^{rew}_{t(i)} = Sensitivity_{rew} * RewardOutcome(t) - Value^{rew}_{t-1(i)}\ if\ i = chosen$$
$$- Value^{rew}_{t-1(i)}\ if\ i = unchosen \quad ..3$$

$$Prediction\ Error^{pun}_{t(i)} = Sensitivity_{pun} * RewardOutcome(t) - Value^{pun}_{t-1(i)}\ if\ i = chosen$$
$$- Value^{pun}_{t-1(i)}\ if\ i = unchosen \quad ..4$$

$$ChoiceProbability_{t(i)} = \frac{exp(Value^{rew}_{t(i)} + Value^{pun}_{t(i)})}{\sum_{j} exp(Value^{rew}_{t(j)} + Value^{pun}_{t(j)})} * (1 - Lapse) + \frac{Lapse}{2} \quad ..5$$

**Model LOOIC comparision:**

Parameters for all models were initially fit for single prior. The winning model was defined as the model with the lowest Leave-One_out Information Criterion (LOOIC):

| Model | Absolute | G:1, L:-1 |
|---|---|---|
| BanditNarm_4par | 17391.53 | 17346.35 |
| **BanditNarm_lapse** | **17207.70** | 17202.98 |
| BanditNarm_lapse_decay | 17223.36 | 17219.81 |
| BanditNarm_singleA_lapse | 17385.94 | 17352.44 |

***Table S3:*** *LOOIC scores for 4 single (full) prior models for absolute reward-punishment and reward-punishment encoded as +1, -1 respectively*

Based on Table S3, we identified Model 2 (banditNarm_lapse) as the winning model based on lowest LOOIC scores.

We then followed up the winning model, with subsequent exploration for different hierarchical priors/ group combinations (Table S4).

| Priors | BanditNarm_lapse |
|---|---|
| Block_conflict_trait (24) | 18446.85 |
| Block_conflict (12) | 18286.42 |
| Block_trait (8) | 17424.22 |
| Conflict_trait (6) | 17636.91 |
| Block (4) | 17405.99 |
| Conflict (3) | 17629.19 |
| Trait (2) | 17212.74 |
| Single prior full (1) | 17207.70 |

***Table S4:*** *LOOIC scores for different priors*

**iii. Control Experiment II**

a. **Game Design:**
Participants completed four experimental blocks/environments based on the varying frequency of Pavlovian congruent and incongruent trials. However, here we report the first two blocks relevant to the present study i.e. Low Conflict Block (LC), High Conflict Block (HC) (see Table S5). Each block comprised 80 trials, equally divided into Go and NoGo.

| Block | Pavlovian Congruent | Pavlovian Incongruent |
|---|---|---|
| Low Conflict | 50% | 50% |
| High Conflict | 30% | 70% |

**Table S5:** Block-design for Experiment II (Orthogonalized Go-NoGo task) showing two subsequent blocks.

In the LC block, trials were evenly split between Pavlovian Congruent (PC) and Pavlovian Incongruent (PI) conditions (50% each) with 25% each of Go and NoGo trials as is the case in a typical Go-NoGo task. We increased the conflict by increasing the frequency of Pavlovian incongruent trials in HC block similar to approach-avoidance conflict task. 30% of trials were Pavlovian Congruent (15 Go and 15 NoGo trials), while 70% were Pavlovian Incongruent (35 Go and 35 NoGo trials). This manipulation was designed to examine whether the asymmetry in motivational conflict that is being required to act or withhold action against Pavlovian tendencies would differentially affect task performance in different environments.

Reward and punishment magnitudes were contingent on action accuracy. For correct responses, outcomes were either a reward of +50 on win trials or no loss (0) on avoid-loss trials, each delivered with an 80% probability. In contrast, incorrect responses resulted in either no reward (0) on win trials or a punishment of −50 on avoid-loss trials, both delivered with 100% probability (Table S6).

| **Action** / **Valence** | **Go** | **NoGo** |
|---|---|---|
| **Win** | 50 / 0 | 50 / 0 |
| **Avoid Loss** | 0 / -50 | 0 / -50 |

**Table S6:** Task outcome contingencies

**b. Data collection procedure:**

83 healthy participants aged 18 years and above were recruited for the study. A total of seven participants were excluded from the final analysis due to technical issues in data collection or storage.

The sequence of the experimental procedure was as follows: Standard Anxiety Test → Instruction Slide → Trial Block → Experimental Blocks → Post Conduction Survey. The experiment was conducted in a controlled laboratory setting with informed consent obtained from all participants. Prior to the task, participants completed the standard questionnaire response for STAI-Trait, STAI-State, GAD7 and PHQ9, comprising multiple subscales assessing various aspects of anxiety and depression symptoms. They were then instructed to read the task guidelines thoroughly and were given the opportunity to practice the trial block as many times as desired to ensure familiarity with the task. Following the practice phase, participants proceeded to the experimental blocks. Upon completion of the task, they were asked to fill out a post-experiment survey, including open-ended questions to gather feedback on their experience, comments, and any criticisms regarding the task or procedure.

For group comparison, the participants were divided based on STAI-Trait scores with median split of the main experiment (M= 43) into High (HTA, n= 41) and Low trait anxiety (LTA, n =35) groups (Figure S5b).

### B. Results:

#### i. Bias construction (supplementary):

**Go Bias:** The average Go bias across participants was 81.3% while average across trait anxiety groups: LTA: 82.3% and HTA: 80.3%.

**Pavlovian Component:** Pavlovian component for No Conflict Tree (NCT) converged near zero for both environments (–0.05 ± 0.50), consistent with the absence of stimulus-driven or color-specific bias. In contrast, HCT elicited robust avoidance (–1.10 ± 0.45) and LCT elicited approach (1.16 ± 0.51). A Kruskal–Wallis test indicated a significant effect of tree type ($\chi^2(2) = 8.41$, $p = 0.015$; Fig. 3b). However, the instrumental component for NCT captures the Pavlovian avoidance bias due to influence of other trees in a particular environment.

**Pavlovian component for HTA and LTA:** Tree type significantly influenced Pavlovian bias in HTA (Kruskal–Wallis $\chi^2(2) = 7.94$, $p = 0.019$), with NCT values converging near zero for both groups (HTA: –0.44 ± 0.81; LTA: 0.25 ± 0.62; Fig. 5b).

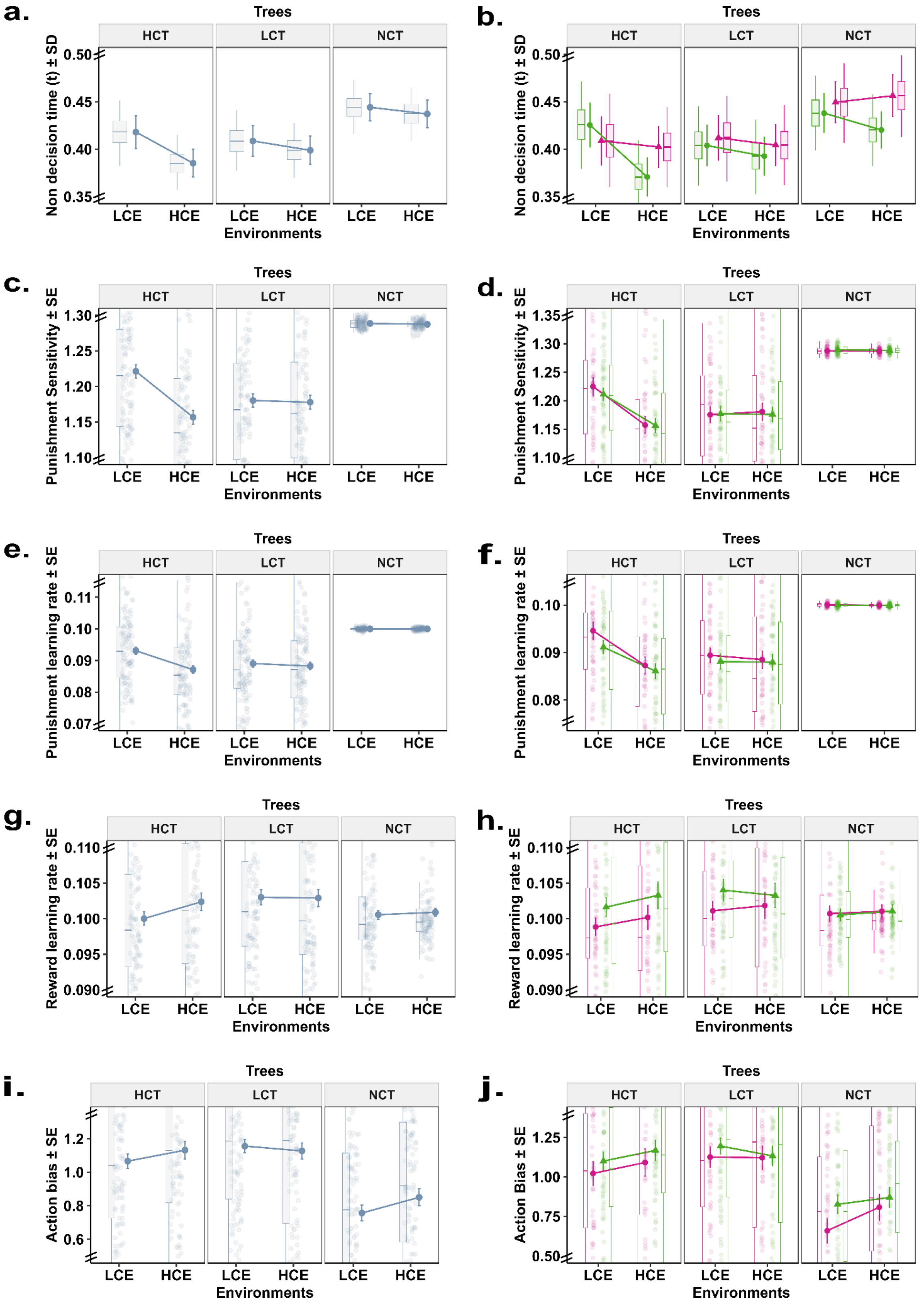


**Figure S1: Additional hierarchical drift–diffusion and reinforcement-learning model parameters**

Mean posterior estimates (± SD) of the non decision time parameter 't' from the Hierarchical Drift Diffusion Model (HDDM) for High, Low, and No conflict trees in LCE and HCE (**a**) and for the two groups LTA and HTA (**b**).

Punishment sensitivity parameter (± SE) from reinforcement learning model in LCE & HCE for High, Low and No conflict trees (**c**) and for the two groups LTA and HTA (**d**).

Punishment learning rate parameter (± SE) from reinforcement learning model in LCE & HCE for High, Low and No conflict trees (**e**) and for the two groups LTA and HTA (**f**).

Reward learning rate parameter (± SE) from reinforcement learning model in LCE & HCE for High, Low and No conflict trees (**g**) and for the two groups LTA and HTA (**h**).

Action bias parameter (± SE) from reinforcement learning model in LCE & HCE for High, Low and No conflict trees (**i**) and for the two groups LTA and HTA (**j**).

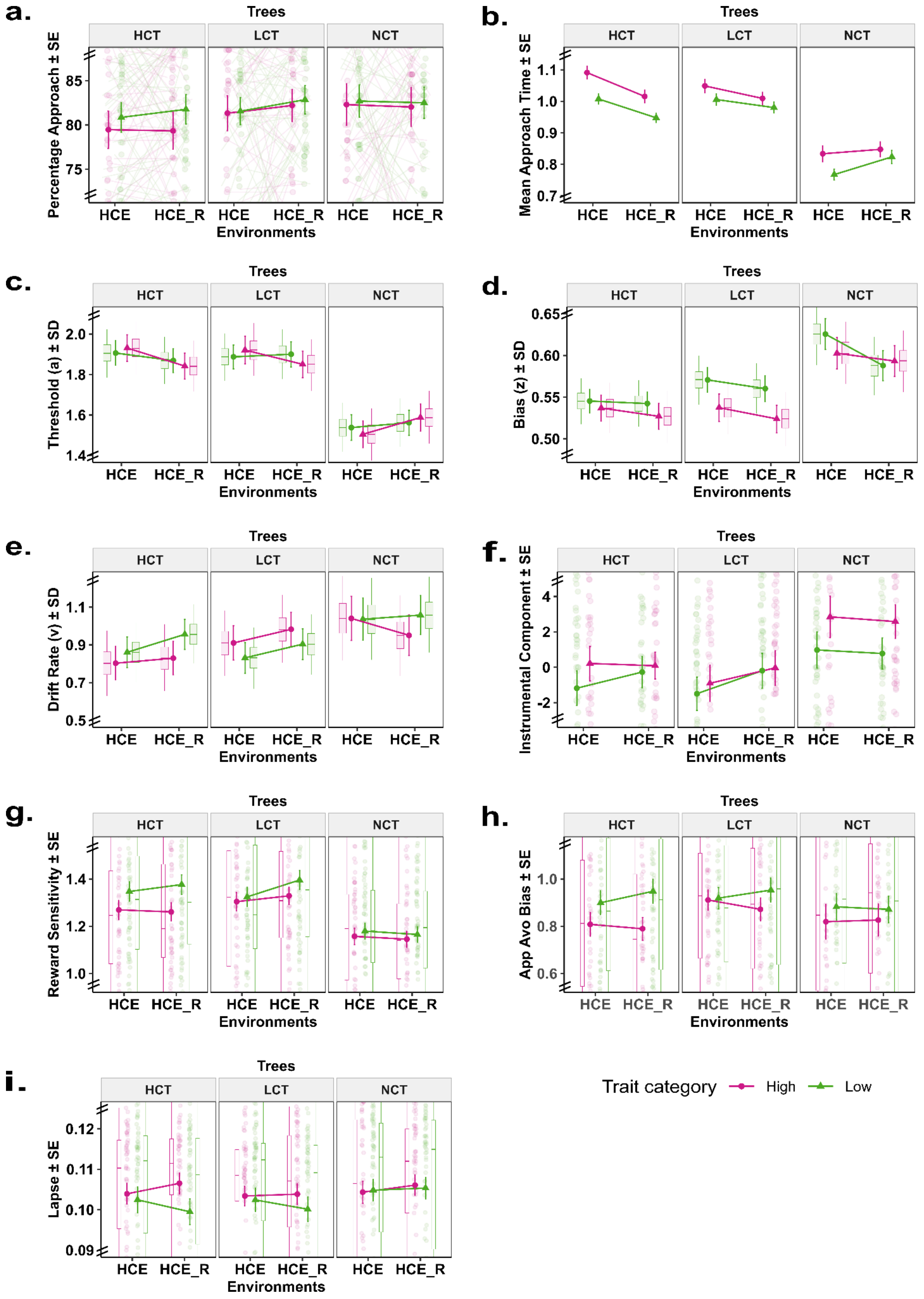


**Figure S2: Behavioural and computational signatures across cue–outcome reversal in a high-conflict environment**

a. Mean percentage approach (± SEM) across HCE and HCE_R for High, Low, and No conflict trees, shown separately for high (pink) and low (green) trait anxiety groups.
b. Mean approach reaction times (in seconds ± SEM) for High, Low, and No conflict stimuli in HCE and HCE_R, shown separately for high (pink) and low (green) trait anxiety groups.
c. Mean posterior estimates (± SD) of the threshold parameter 'a' from the Hierarchical Drift Diffusion Model (HDDM) for High, Low, and No conflict stimuli in HCE and HCE_R, shown separately for high (pink) and low (green) trait anxiety groups.
d. Mean posterior estimates (± SD) of the starting bias parameter 'z' from the Hierarchical Drift Diffusion Model (HDDM) for High, Low, and No conflict stimuli in HCE and HCE_R, shown separately for high (pink) and low (green) trait anxiety groups.
e. Mean posterior estimates (± SD) of the drift rate parameter 'v' from the Hierarchical Drift Diffusion Model (HDDM) for High, Low, and No conflict stimuli in HCE and HCE_R, shown separately for high (pink) and low (green) trait anxiety groups.
f. Instrumental component with SE, for High, Low and No conflict trees for HCE and HCE_R, shown separately for high (pink) and low (green) trait anxiety groups.
g. Reward sensitivity parameter with SE from reinforcement learning model in HCE and HCE_R for High, Low and No conflict trees, shown separately for high (pink) and low (green) trait anxiety groups.
h. Approach-avoid bias parameter with SE from reinforcement learning model in HCE and HCE_R for High, Low and No conflict trees, shown separately for high (pink) and low (green) trait anxiety groups.
i. Lapse parameter with SE from reinforcement learning model in HCE and HCE_R for High, Low and No conflict trees, shown separately for high (pink) and low (green) trait anxiety groups.

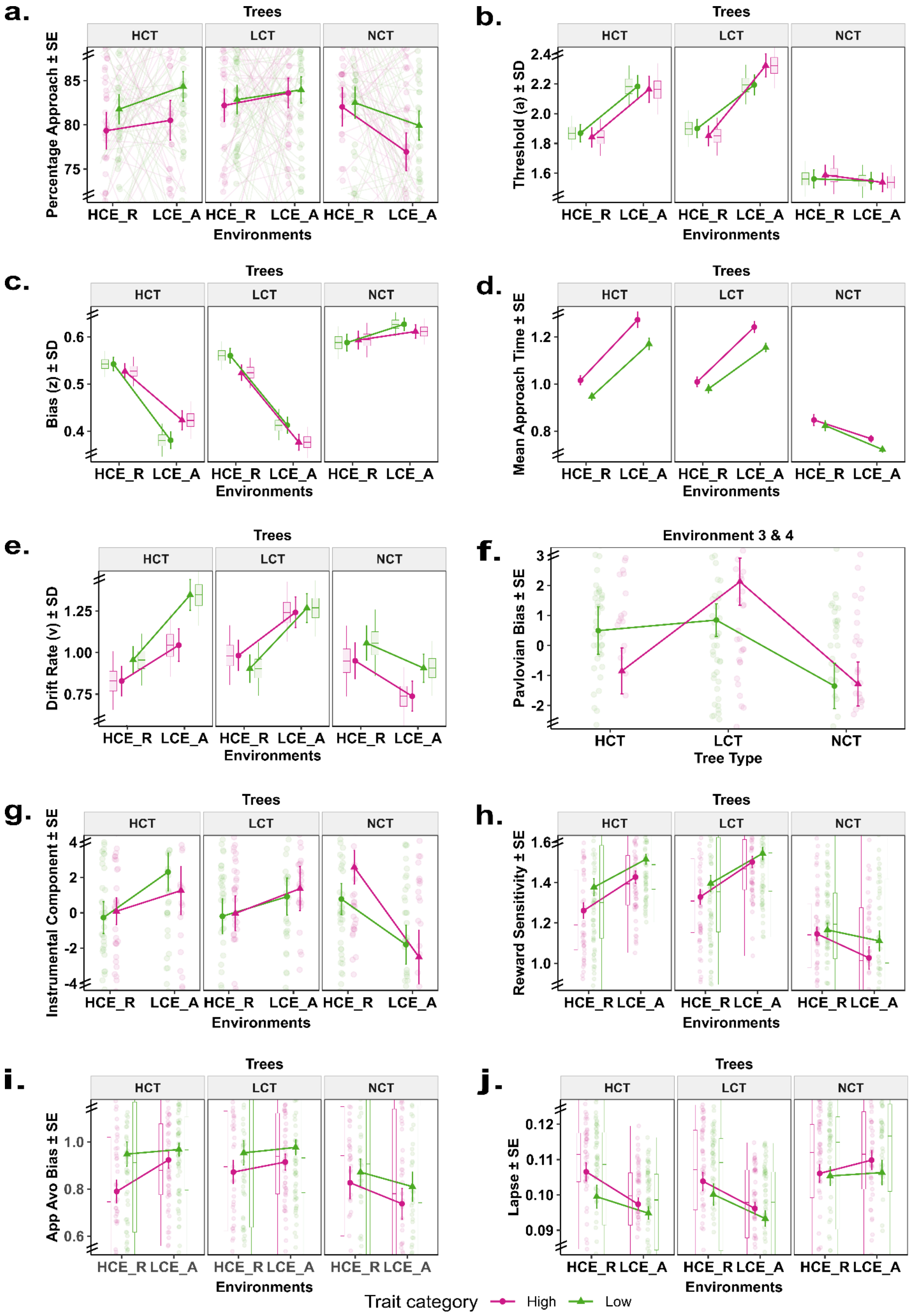


**Figure S3: Behavioural and computational patterns in a low-conflict but aversive environment**

a. Mean percentage approach (± SEM) across HCE_R and LCE_A for High, Low, and No conflict trees, shown separately for high (pink) and low (green) trait anxiety groups.
b. Mean posterior estimates (± SD) of the threshold parameter 'a' from the Hierarchical Drift Diffusion Model (HDDM) for High, Low, and No conflict stimuli in HCE_R and LCE_A, shown separately for high (pink) and low (green) trait anxiety groups.
c. Mean posterior estimates (± SD) of the starting bias parameter 'z' from the Hierarchical Drift Diffusion Model (HDDM) for High, Low, and No conflict stimuli in HCE_R and LCE_A, shown separately for high (pink) and low (green) trait anxiety groups.
d. Mean approach reaction times (in seconds ± SEM) for High, Low, and No conflict stimuli in HCE_R and LCE_A, shown separately for high (pink) and low (green) trait anxiety groups.
e. Pavlovian bias with SEM, for High, Low and No conflict trees for Environment 3-4, shown separately for high (pink) and low (green) trait anxiety groups.
f. Mean posterior estimates (± SD) of the drift rate parameter 'v' from the Hierarchical Drift Diffusion Model (HDDM) for High, Low, and No conflict stimuli in HCE_R and LCE_A, shown separately for high (pink) and low (green) trait anxiety groups.
g. Instrumental component with SEM, for High, Low and No conflict trees for HCE_R and LCE_A, shown separately for high (pink) and low (green) trait anxiety groups.
h. Reward sensitivity parameter with SEM from reinforcement learning model in HCE_R and LCE_A for High, Low and No conflict trees, shown separately for high (pink) and low (green) trait anxiety groups.
i. Approach-avoid bias parameter with SEM from reinforcement learning model in HCE_R and LCE_A for High, Low and No conflict trees, shown separately for high (pink) and low (green) trait anxiety groups.
j. Lapse parameter with SEM from reinforcement learning model in HCE_R and LCE_A for High, Low and No conflict trees, shown separately for high (pink) and low (green) trait anxiety groups.

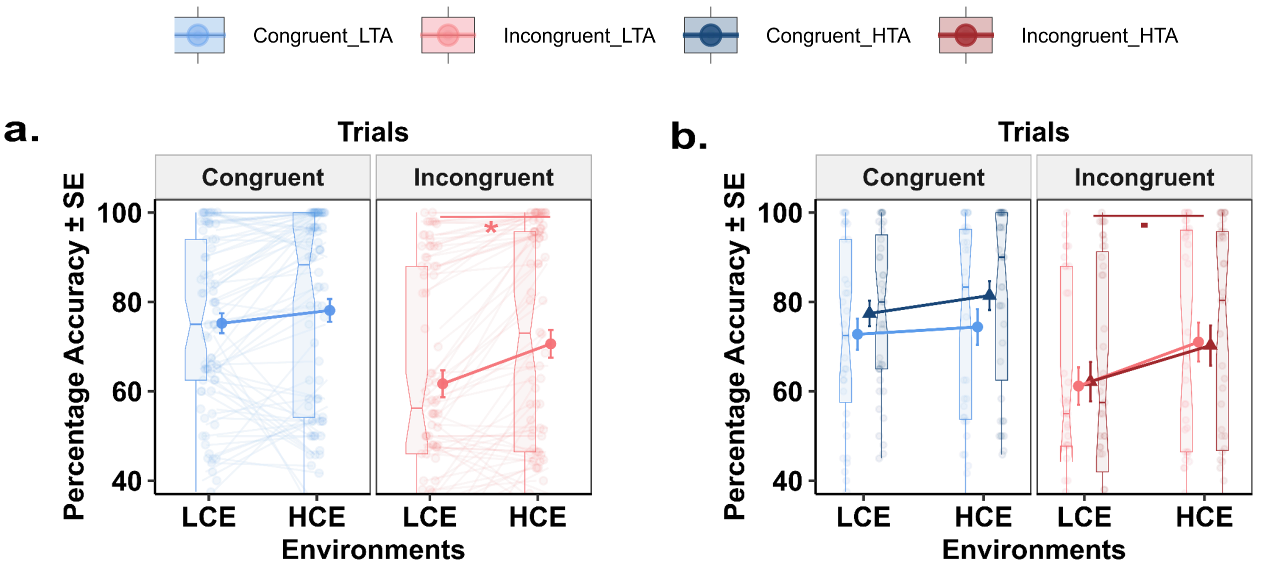


**Figure S4: Effects of environmental conflict on Go/NoGo task performance**

a. Percentage accuracy (± SEM) across LCE and HCE for the two trial types Pavlovian congruent (blue) and Pavlovian incongruent (red) in orthogonalized Go-NoGo task.

**b.** Percentage accuracy (± SEM) across LCE and HCE for the two trial types Pavlovian congruent and incongruent for two trait anxiety groups HTA (dark blue and maroon) and LTA (light blue and light red) in orthogonalized Go-NoGo task.

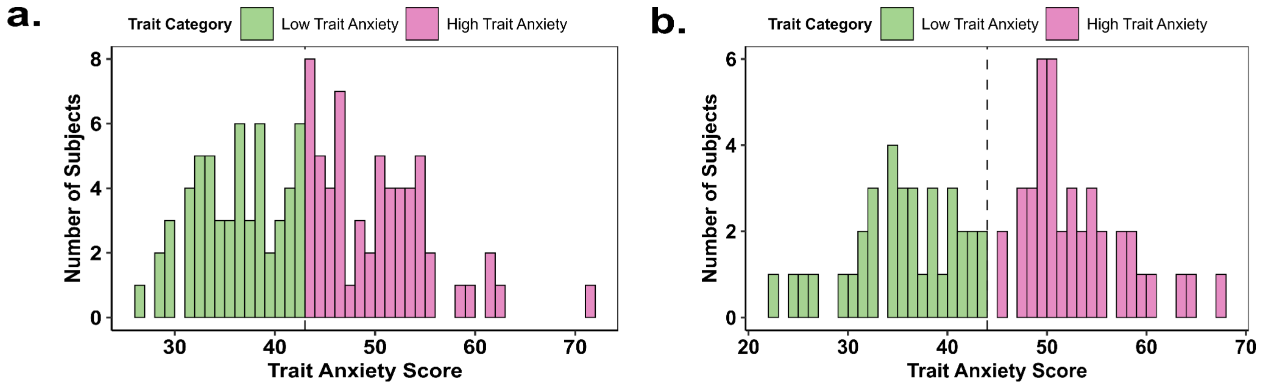


**Figure S5: Distribution of trait anxiety scores with median-based grouping across experimental samples**

a. Histogram showing distribution of Trait anxiety scores in the Experiment I (Approach avoidance conflict paradigm) sample also median split = 43 division into Trait category : LTA (green) and HTA (pink).

b. Histogram showing distribution of Trait anxiety scores in the Experiment II (Go-NoGo paradigm) sample also median split = 43 division into Trait category : LTA (green) and HTA (pink).
